# ESCPE-1 Mediates Retrograde Endosomal Sorting of the SARS-CoV-2 Host Factor Neuropilin-1

**DOI:** 10.1101/2022.01.20.477115

**Authors:** Boris Simonetti, James L. Daly, Lorena Simón-Gracia, Katja Klein, Saroja Weeratunga, Carlos Antón-Plágaro, Allan Tobi, Lorna Hodgson, Phil Lewis, Kate J. Heesom, Deborah K. Shoemark, Andrew D. Davidson, Brett M. Collins, Tambet Teesalu, Yohei Yamauchi, Peter J. Cullen

**Author notes:** Corresponding author (B.S.); (P.J.C.). These authors contributed equally to this work (Boris Simonetti, James L. Daly).

## Abstract

Endosomal sorting maintains cellular homeostasis by recycling transmembrane proteins and associated proteins and lipids (termed ‘cargoes’) from the endosomal network to multiple subcellular destinations, including retrograde traffic to the *trans*-Golgi network (TGN). Viral and bacterial pathogens subvert retrograde trafficking machinery to facilitate infectivity. Here, we develop a proteomic screen to identify novel retrograde cargo proteins of the Endosomal SNX-BAR Sorting Complex Promoting Exit-1 (ESCPE-1). Using this methodology, we identify Neuropilin-1 (NRP1), a recently characterised host factor for severe acute respiratory syndrome coronavirus 2 (SARS-CoV-2) infection, as a cargo directly bound and trafficked by ESCPE-1. ESCPE-1 mediates retrograde trafficking of engineered nanoparticles functionalised with the NRP1-interacting peptide of the SARS-CoV-2 Spike protein. ESCPE-1 sorting of NRP1 may therefore play a role in the intracellular membrane trafficking of NRP1-interacting viruses such as SARS-CoV-2.

---

Retrograde transport from endosomes to the *trans*-Golgi network (TGN) diverts integral proteins away from lysosomal degradation to facilitate a variety of cellular functions^1^, and is exploited by multiple pathogens^2–8^. Despite technical advances into the study of this process^9–13^, the broad array of cargoes that traverse this pathway and the mechanistic basis of their sorting largely remains elusive due to the lack of tools for the unbiased interrogation of the dynamic TGN proteome. To overcome this, we engineered a peroxidase-based proximity biotinylation methodology by fusing horseradish peroxidase (HRP) to the luminal terminus of TGN46, a representative single-pass type I transmembrane protein that predominantly localises to the TGN^14^. Following incubation with the membrane-permeable precursor biotin-phenol (BP) and a brief 60 second addition of hydrogen peroxide (H_2_O_2_), HRP generates short-lived, membrane-impermeable biotin-phenoxyl radicals that irreversibly biotinylate vicinal soluble proteins and the luminal domains of transmembrane proteins (**Fig. 1A**)^15^.

**Fig. 1.**
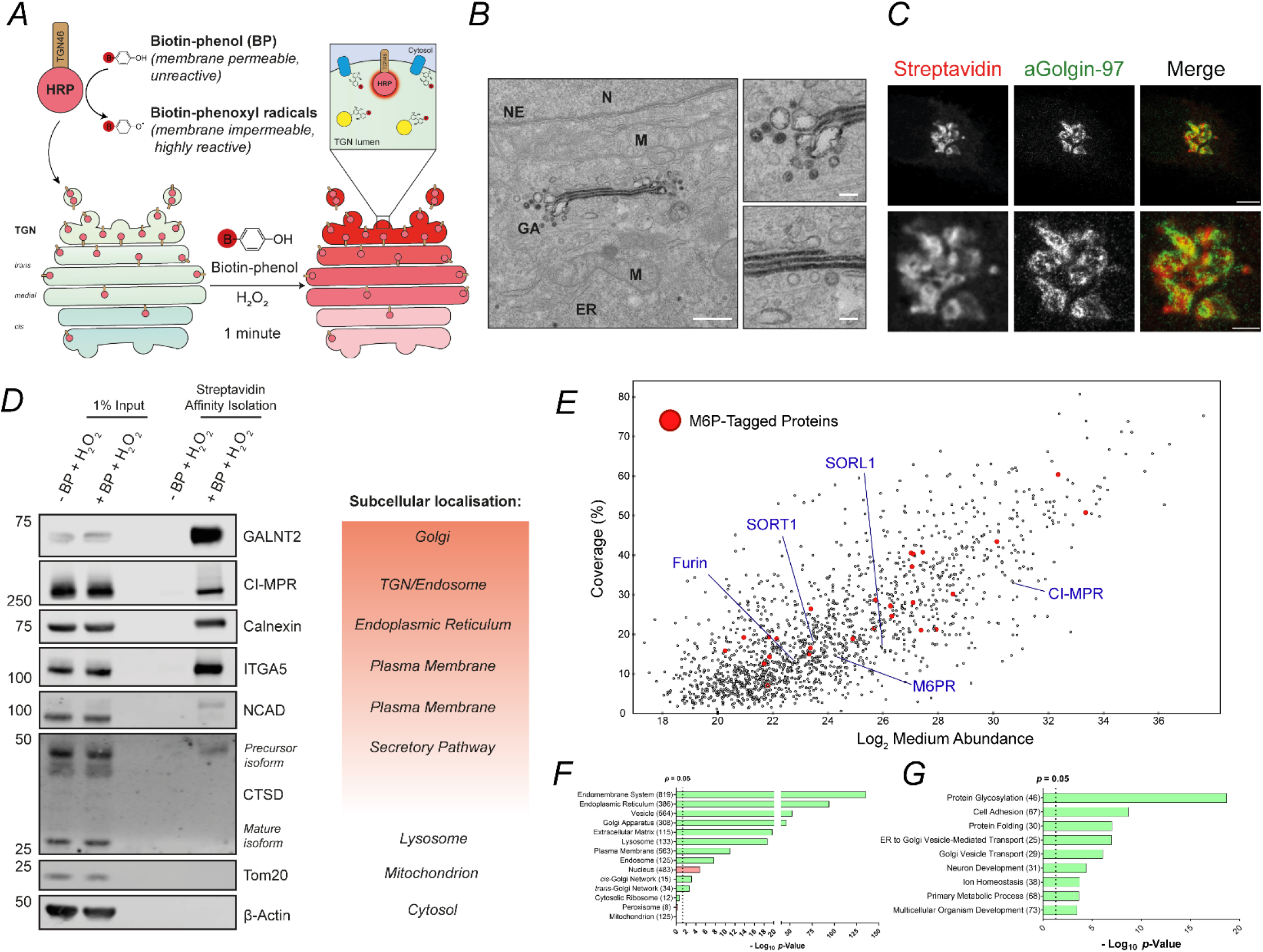
Development of a Methodology to Biotinylate TGN-Resident Proteins. **(A)** Schematic of HRP-TGN46 biotinylation labelling of endogenous TGN-resident proteins. In the presence of biotin-phenol and hydrogen peroxide, HRP catalyses the formation of membrane-impermeable biotin-phenoxyl that covalently label nearby endogenous proteins. **(B)** Transmission electron micrograph of HRP-TGN46-expressing HeLa cells incubated with DAB and H_2_O_2_ for 10 minutes. Scale bar: 500 nm, zoom scale bar: 100 nm. ER = Endoplasmic Reticulum, GA = Golgi Apparatus, M = Mitochondrion, N = Nucleus, NE = Nuclear Envelope. **(C)** Stimulated emission depletion (STED) microscopy of HRP-TGN46-expressing HeLa cells following biotinylation, stained with fluorescent streptavidin and immunofluorescence labelling of Golgin-97. Scale bars = 5 µm, 2 µm insets. **(D)** Streptavidin affinity isolation of biotinylated proteins from total cell lysate of HRP-TGN46-expressing cells incubated with H_2_O_2_ in the presence or absence of BP, and their corresponding known subcellular localisation. **(E)** Scatter plot of all HRP-TGN46 labelled proteins, with known mannose-6-phosphate-tagged proteins overlaid in red, and example TGN retrograde cargoes highlighted by blue labels. N = 5 independent repeats. **(F-G)** Graphical representation of overrepresentation of gene ontology cellular compartments (F) and biological processes (G) within the HRP-TGN46-proteome according to their calculated p-value. Overrepresented gene ontology terms are coloured green, and underrepresented terms are coloured red. Brackets represent the number of identified proteins belonging to each category.

In a HeLa cell line stably expressing HRP-TGN46 (**Extended Data Fig. 1A**), biotinylation was exclusively observed upon the presence of HRP-TGN46, BP and H_2_O_2_ (**Extended Data Fig. 1B**). By either light microscopy, visualised by fluorescent streptavidin staining, or electron microscopy, visualised by oxidative polymerisation of the electron-dense 3,3’-diaminobenzidine (DAB), we observed predominant labelling of the TGN upon activation of HRP-TGN46 (**Fig. 1B,C** **and Extended Data Fig. 1C,D**). The streptavidin-based affinity isolation of whole cell lysates following biotinylation showed specific labelling of endogenous proteins spanning the secretory pathway, including the Golgi-localised precursor isoform of the lysosomal hydrolase cathepsin D (CTSD); the glycosylation enzyme polypeptide N-acetylgalactosaminyltransferase 2 (GALNT2); the retrograde TGN-resident cargo cation-independent mannose 6-phosphate receptor (CI-MPR); the endoplasmic reticulum (ER) protein calnexin; and cell surface receptors such as integrin-α5 (ITGA5) and N-cadherin (NCAD), that traverse the biosynthetic pathway *en route* to the plasma membrane (**Fig. 1D**). SILAC-based mass spectrometry revealed a list of 1237 proteins reproducibly enriched by streptavidin affinity isolation following HRP-TGN46 labelling (**Fig. 1E, Extended Data Fig. 1E,F Supplementary Tables 1,2**). Gene ontology analysis demonstrated an enrichment of cellular components corresponding to the biosynthetic pathway and the interface of the TGN with the endolysosomal network, including a 74% coverage of TGN-resident mannose-6-phoshphate tagged proteins^17^, and validated retrograde transmembrane cargoes including Furin, CI-MPR, M6PR, SORT1 and SORL1 (**Fig. 1F,G, Supplementary Tables 3-5**).

ESCPE-1 is an endosomal coat complex implicated in the retrograde trafficking of transmembrane proteins such as the CI-MPR^13,18–20^. ESCPE-1 consists of a heterodimer of SNX1 or SNX2, which are functionally redundant, associated with either SNX5 or SNX6, which are also functionally redundant and mediate direct binding to the cytosolic tails of transmembrane cargo^18–21^ (**Fig. 2A**). Recent advances have revealed the identity of transmembrane cargoes that undergo sequence-dependent recognition by ESCPE-1 for sorting to the plasma membrane^20,22^, yet comparatively less is known about cargoes re-routed towards the TGN.

**Fig. 2.**
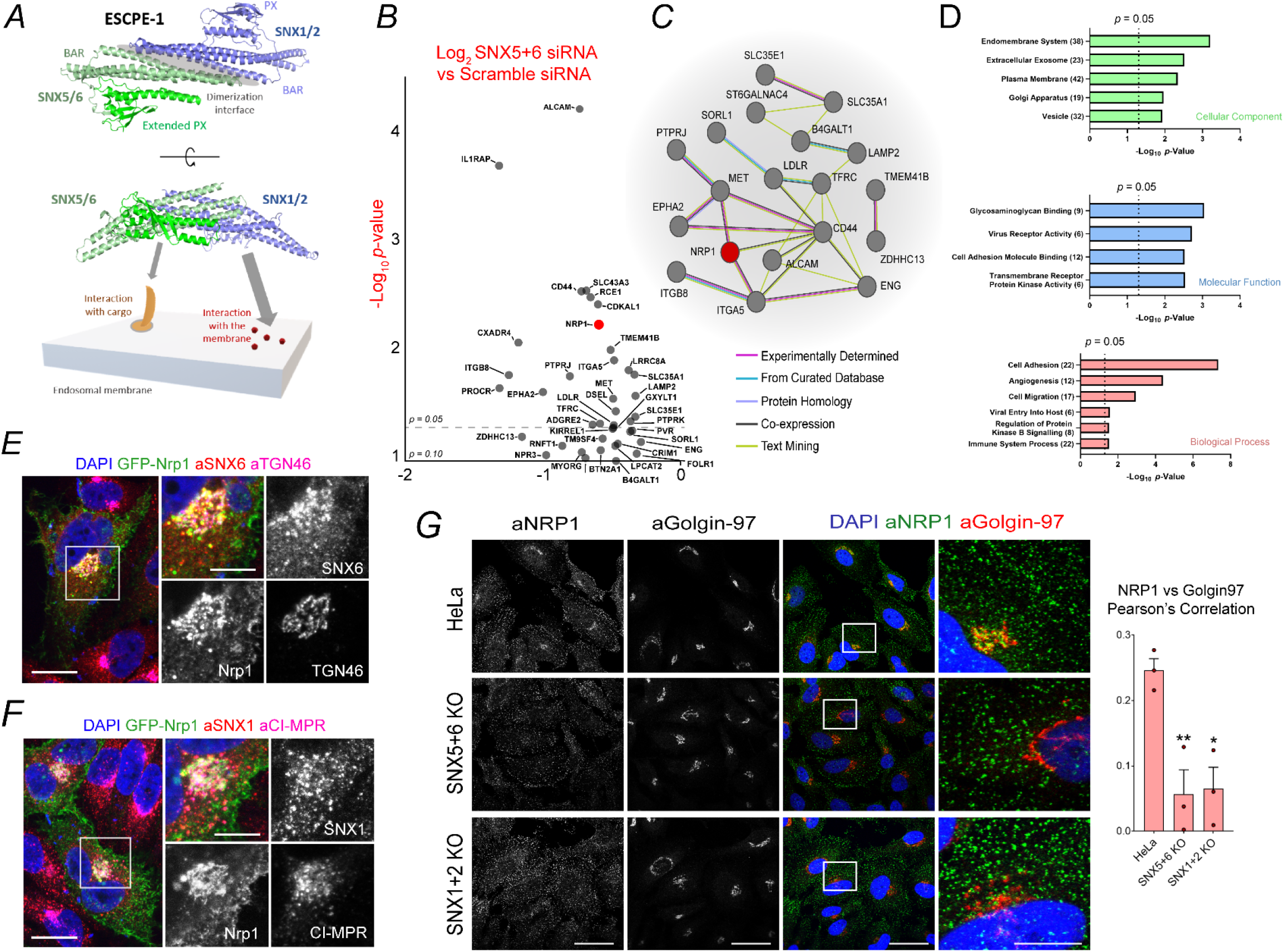
A Proteomic Screen for ESCPE-1 Retrograde Cargoes Identifies NRP1. **(A)** Schematic of ESCPE-1 dimerisation and coincidence detection of endosomal membranes and transmembrane cargo. **(B)** Volcano plot of transmembrane proteins depleted from the HRP-TGN46-labelled proteome upon SNX5+SNX6 suppression by < Log_2_ -0.26, p value < 0.1 by SILAC proteomics. p = 0.05 is indicated by a dotted line. NRP1 is highlighted in red. N = 4 independent repeats **(C)** STRING network analysis of the transmembrane proteins presented in (B). The legend indicates the level of evidence for each protein-protein association. **(D)** Gene ontology analysis of the transmembrane proteins identified in (B), classified by cellular component, molecular function and biological process. Graphs represent the statistical significance of category enrichment, with a dotted line representing p = 0.05. Brackets represent the number of proteins in each category. **(E-F)** HeLa cells expressing GFP-Nrp1 were co-stained with anti-SNX6 and anti-TGN46 antibodies (E), or anti-SNX1 and anti-CI-MPR antibodies (F). Scale = 20 µm, zoom = 10 µm. **(G)** Immunofluorescence staining of endogenous NRP1 in HeLa WT cells, SNX5+6 KO HeLa cells and SNX1+2 KO HeLa cells. Scale = 50 µm, insets 5 µm. Pearson’s correlation quantification of the colocalisation of NRP1 and Golgin-97 upon ESCPE-1 depletion Ordinary one-way ANOVA with Dunnett’s multiple comparison’s tests. N = 3 independent repeats. WT vs SNX5+6 KO p = 0.0084, WT vs SNX1+2 KO p = 0.0103.

To identify the transmembrane proteins that undergo ESCPE-1-dependent retrograde transport, SILAC-based quantitative proteomics was used to compare HRP-TGN46-mediated biotinylation in Scramble (Scr) siRNA-treated and double SNX5+SNX6 siRNA-treated HeLa cells (due to functional redundancy both SNX5 and SNX6 must be suppressed to perturb ESCPE-1-dependent trafficking)^18,19^. Retrograde trafficking of TGN46 has been demonstrated to be independent of ESCPE-1^18,19^. Double SNX5+SNX6 knockdown did not perturb HRP-TGN46 labelling at the whole cell lysate level (**Extended Data Fig. 2A**).

Of the previously established list of proteins biotinylated by HRP-TGN46, 46 proteins were significantly depleted (p < 0.05 Log_2_ fold-change < -0.26) in the SNX5+SNX6 siRNA HRP-TGN46-labelled proteome compared to Scramble siRNA-treated cells, of which 25 contained transmembrane-spanning domains (**Fig. 2B, Extended Data Fig. 2B and Supplementary Tables 6-8**). STRING analysis identified a network of protein-protein interactions, including a cluster of proteins containing Neuropilin-1 (NRP1) alongside integrin-α5, integrin-β8, CD44 and the receptor tyrosine kinases MET and EPHA2 (**Fig. 1C**). Biological processes and molecular function categories pertaining to cellular adhesion and migration, transmembrane receptor kinase activity, and virus receptor activity were significantly enriched (**Fig. 2D, Supplementary Tables 9-11**). Moreover, comparison of these proteins with published ESCPE-1 interactors and cell surface cargoes revealed a consistent enrichment of biological pathways (**Extended Data Fig. 2C**).

NRP1 is a co-receptor for a range of extracellular ligands, including members of the vascular endothelial growth factor (VEGF) and semaphorin families, and was recently identified as a host factor that facilitates SARS-CoV-2 infection^23,24^. Moreover, NRP1 was also enriched in previously published interaction networks of SNX5, SNX6 (and the neuronal SNX6 paralogue SNX32), though its surface levels were unaffected by ESCPE-1 inactivating mutations^19,20^ (**Extended Data Fig. 2D**). Expression of an extracellular/luminally GFP-tagged murine NRP1 (GFP-Nrp1) revealed predominant localisation to the plasma membrane and an internal population that colocalised with SNX1- and SNX6-decorated endosomes, and with the TGN markers TGN46, Golgin-97 and CI-MPR (**Fig. 2E,F**). Moreover, we confirmed by immunofluorescence staining that the TGN-resident pool of endogenous NRP1 was reduced in HeLa cells lacking ESCPE-1 subunits (**Fig. 2G**).

The extracellular/luminal GFP-tag on the Nrp1 construct was amenable for an antibody uptake assay to facilitate the chase of internalised GFP-Nrp1 from the cell surface into intracellular compartments following endocytosis (**Extended Data Fig. 3A**). At early timepoints GFP-Nrp1 colocalised with intracellular vesicles, a sub-population of which were SNX6-positive endosomes, and progressively colocalised with TGN46 at later timepoints (**Fig. 3A, Extended Data Fig. 3B**). 3D reconstruction of the TGN46 signal revealed a population of internalised GFP-Nrp1 within the TGN (**Extended Data Fig. 3C**).

**Fig. 3.**
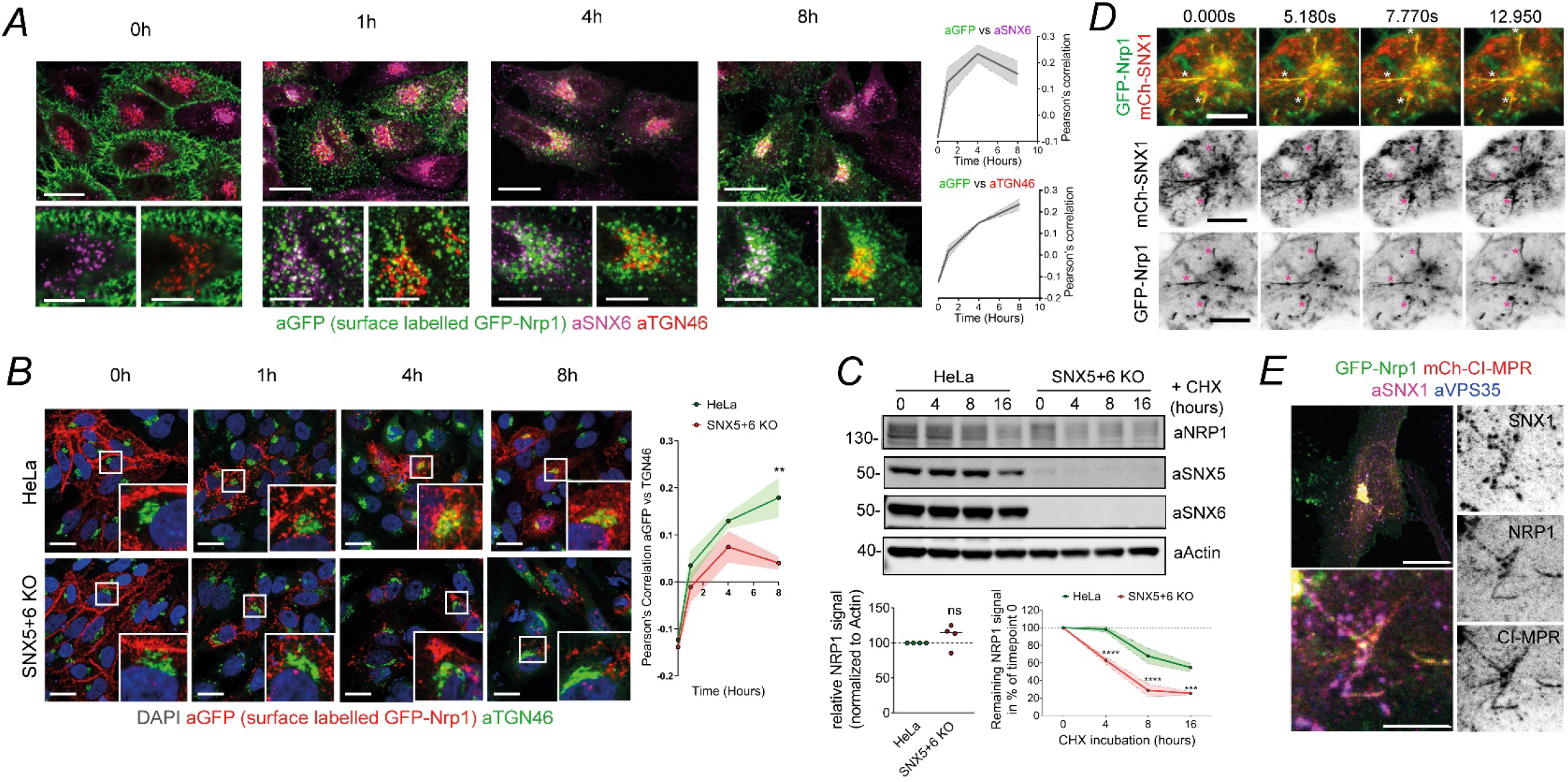
ESCPE-1 Mediates Tubular Retrograde Trafficking of NRP1. **(A)** Anti-GFP antibodies were bound to extracellular-facing GFP-Nrp1 in HeLa, followed by a surface uptake assay. At 0, 1, 4 and 8 hour timepoints, cells were fixed and labelled with anti-SNX6 and anti-TGN46 antibodies. N = 3 independent repeats. **(B)** GFP-Nrp1 surface uptake assay in wild-type HeLa cells or SNX5+6 KO HeLa cells. Pearson’s correlation between anti-GFP and anti-TGN46 is measured over time. Colocalisation was measured over N = 4 experiments with a two-way ANOVA and Šídák’s multiple comparisons test, 0h: p = 0.9897, 1h: p = 0.6504, 4h: p = 0.4805, 8h: p = 0.0041. Scale = 20 µm**. (C)** Degradation assay in HeLa and HeLa SNX5+SNX6 KO cells. Cells were incubated with 10 µg/ml cycloheximide and lysed at different time points as indicated. The band intensity of endogenous NRP1 was measured from N = 4 independent experiments using Odyssey software and was normalised to wild-type HeLa levels. Steady state levels of NRP1 were comparable between HeLa and SNX5+6 KO cells (left graph), Two-tailed unpaired t test; p = 0.2803. The levels of NRP1 at different timepoints were compared using a two-way ANOVA and Šídák’s multiple comparisons test (right graph). 4 hour: p < 0.0001, 8 hour: p = < 0.0001, 16 hour : p = 0.0008. **(D)** HeLa cells were cotransfected with GFP-Nrp1 and mCherry-SNX1 and live imaged after 24 hours. Asterisk = examples of GFP-Nrp1 and SNX1 signals colocalising in tubular profiles emanating from endosomes. Scale bar = 10 µm. **(E)** Confocal microscopy of HeLa cells co-transfected with mCherry-CI-MPR and GFP-Nrp1, co-stained with anti-SNX1 and anti-VPS35 antibodies. Scale = 20 µm, zoom = 5 µm. The bars, error bars and circles represent the mean, SEM and individual data points, respectively. ∗p< 0.05, ∗∗p< 0.01, ∗∗∗p< 0.001, ∗∗∗∗p< 0.0001.

When we repeated the uptake assay in a previously described SNX5+SNX6 double KO HeLa cell line^20^, GFP-Nrp1 showed a decreased rate of colocalisation with TGN46 (**Fig. 3B**). Blocking of protein synthesis with cycloheximide (CHX) revealed an increased rate of endogenous NRP1 turnover in SNX5+SNX6 KO cells, consistent with enhanced lysosomal degradation in the absence of retrograde endosomal sorting (**Fig. 3C**). Importantly no significant change of GFP-Nrp1 trafficking was observed at earlier time-points, suggesting that internalisation and early endocytic trafficking of GFP-Nrp1 is not altered in the SNX5+SNX6 KO cells (**Fig. 3B**).

ESCPE-1 mediates retrograde endosomal trafficking by driving the biogenesis of cargo-enriched tubulovesicular membrane carriers that emanate from endosomes and couple to dynein/dynactin motor complexes for transport towards the TGN^1,25–27^. Live imaging of cells co-expressing GFP-Nrp1 and mCherry-SNX1 revealed enrichment of the receptor in tubular profiles emanating from SNX1 endosomes (**Fig. 3D, Extended Data Fig. 3D, Supplementary Videos 1,2**). To assess whether these were indeed tubular carriers undergoing endosome-to-TGN trafficking we co-expressed GFP-Nrp1 alongside a mCherry-CI-MPR construct, which is the prototypical ESCPE-1 retrograde cargo^18–20^. Fluorescently tagged Nrp1 and CI-MPR colocalised in the same tubular profiles (**Fig. 3E, Extended Data Fig. 3E**, **Supplementary Video 3**), and endogenous SNX1 was found to decorate portions of these transport carriers (**Fig. 3E**). We conclude that ESCPE-1 mediates tubular-based sorting of NRP1 from endosomes to the TGN, through a similar pathway to that of CI-MPR.

GFP-nanotrap of GFP-tagged ESCPE-1 subunits revealed that SNX5, SNX6 and SNX32 were able to immunoprecipitate endogenous NRP1, alongside CI-MPR (**Fig. 4A**). Consistently, NRP1 was also enriched in SILAC-based interactomes of SNX5, SNX6 and SNX32^19^ (**Extended Data Fig. 4A**). The interaction with SNX5 and SNX6 was also observed for the NRP1 homologue NRP2 (**Fig. 4B, Extended Data Fig. 4B,C**). Conversely, GFP-tagged NRP1 tail immunoprecipitated mCherry-tagged SNX5 and SNX6, and all endogenous ESCPE-1 subunits (**Extended Data Fig. 4D,E**).

**Fig. 4.**
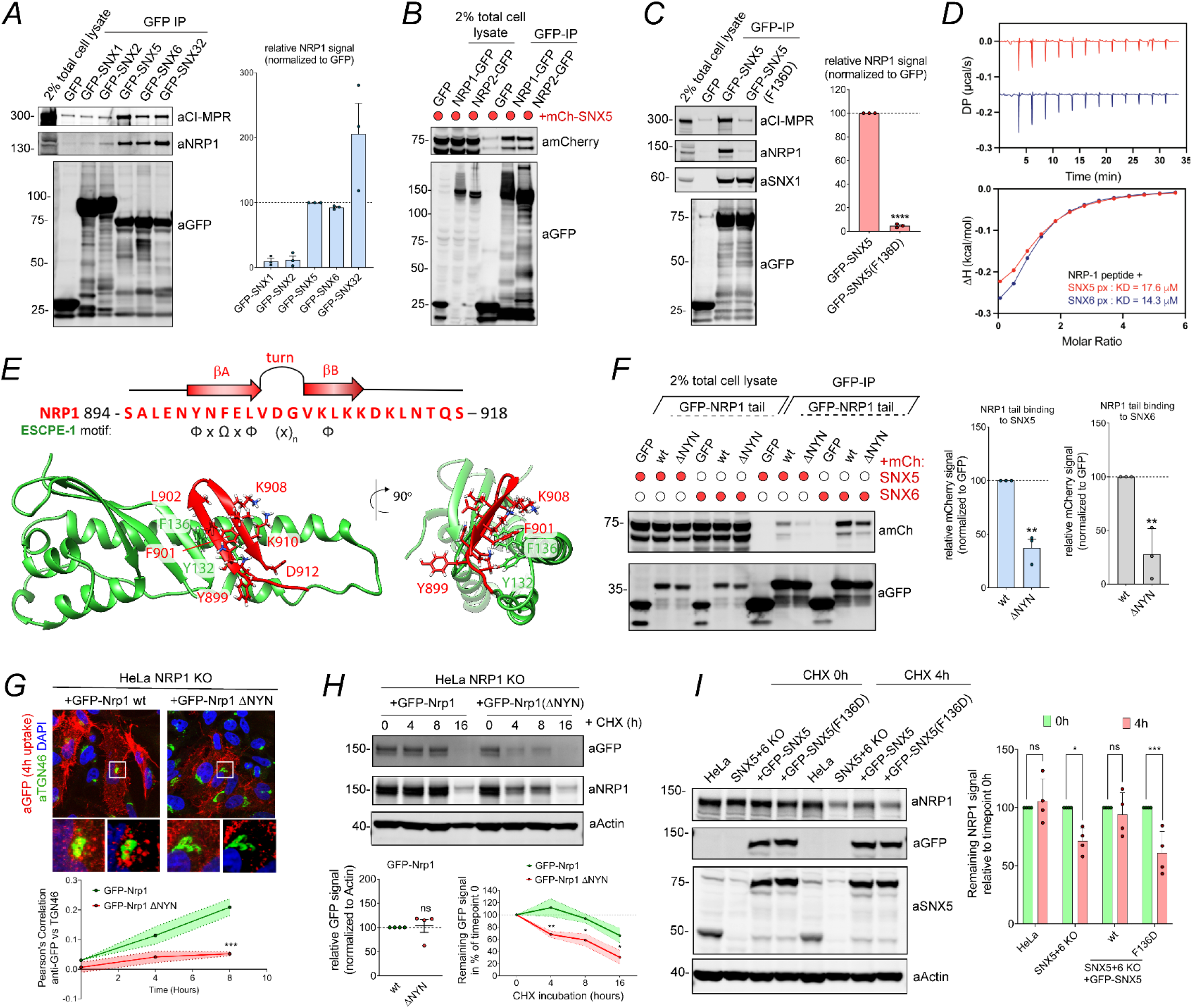
Molecular Basis for the ESCPE-1 Interaction with NRP1. **(A)** HEK293T cells were transfected to express GFP-tagged SNX1, SNXf2, SNX5, SNX6 and SNX32 and subjected to GFP-nanotrap (N = 3 independent experiments). **(B)** HEK293T cells were cotransfected to express GFP-tagged NRP1 or NRP2 and mCherry or mCherry-tagged SNX5 and subjected to GFP-nanotrap. The blot is representative of three independent experiments. **(C)** HEK293T cells were transfected to express GFP-tagged SNX5 or SNX5 F136D mutant and subjected to GFP-nanotrap (N = 3 independent experiments). Two-tailed unpaired t test; p <0.0001. **(D)** SNX5 and SNX6 PX domain were titrated against the NRP1 peptide and binding was measured by ITC. Top panel shows the raw data and bottom panel shows the integrated and normalised data fitted with a 1:1 binding model. The kD values were measured over N = 3. **(E)** Top: schematic of the NRP1 cytosolic domain sequence highlighting the residues that conform to the ESCPE-1 binding motif and that are predicted to fold into the beta-hairpin structure that engages the PX domain of SNX5. Bottom: molecular model of the cytosolic tail of NRP1 bound to the extended PX domain of SNX5. **(F)** HEK293T cells were cotransfected to express GFP-tagged NRP1 tail wild type or a ΔNYN mutant and mCherry-tagged SNX5 or SNX6 and subjected to GFP-nanotrap. SNX5 binding: Two-tailed unpaired t test; p = 0.0013. SNX6 binding: Two-tailed unpaired t test; p = 0.0064. (**G**) HeLaNRP1KO cells transfected with GFP-Nrp1 or GFP-Nrp1-ΔNYN mutant constructs were subjected to an antibody uptake assay for 0, 4 or 8 hours. At the different timepoints cells were fixed and stained for TGN46, and anti-GFP colocalisation was quantified in the GFP-positive subpopulation of cells (N = 3 independent experiments). Two-way ANOVA and Šídák’s multiple comparisons test, 0 hour p = 0.8060, 4 hours p = 0.0736, 8 hours p = 0.0004. **(H)** Degradation assay in HeLaNRP1KO cells transiently transfected with GFP-Nrp1 or GFP-Nrp1 ΔNYN mutant. 48h after transfection, cells were incubated with 10 µg/ml cycloheximide and lysed at different time points as indicated. The band intensity of GFP through was measured from N = 4 independent experiments using Odyssey software. At steady state,GFP-Nrp1 or GFP-Nrp1 ΔNYN constructs expressed at comparable levels (left graph), two-tailed unpaired t test; p = 0.8147. The levels of GFP-Nrp1 or GFP-Nrp1 ΔNYN at different timepoints were compared using a two-way ANOVA and Šídák’s multiple comparisons test (right graph). 4h: p = 0.0091, 8h: p = 0.0438, 16h: p = 0.0397. **(I)** Degradation assay in HeLa, HeLa SNX5+6KO transduced to express GFP-SNX5 or the GFP-SNX5(F136D) mutant. Cells were incubated with 10 µg/ml cycloheximide and lysed at different time points as indicated. The band intensity of endogenous NRP1 was measured from N = 4 independent experiments using Odyssey software. In each cell line the degradation of NRP1 after 4h was compared to the 0h levels using a two-way ANOVA and Šídák’s multiple comparisons test. HeLa: p = 0.9455; SNX5+6 KO: p = 0.0112; +GFP-SNX5: p = 0.9416; +GFP-SNX5(F136D): p = 0.0005. The bars, error bars and circles represent the mean, SEM and individual data points, respectively. ∗p< 0.05, ∗∗p< 0.01, ∗∗∗p< 0.001, ∗∗∗∗p< 0.0001.

ESCPE-1 cargoes possess a ФxΩxФx_n_Ф sorting motif in their cytosolic tail (whereby Ф corresponds to hydrophobic residues and Ω represents a central aromatic residue) that folds into a β-hairpin structure (comprised of two strands, denoted βA and βB, interspaced by a flexible loop of variable length, denoted x_n_) that is recognised by the extended PX domain of SNX5 and SNX6^20,28^. We mapped NRP1 binding to the SNX5 PX domain by engineering a SNX1 chimeric construct where the SNX1 PX domain was replaced with that of SNX5; this successfully immunoprecipitated endogenous NRP1 (**Extended Data Fig. 4F**). Mutagenesis of the F136 residue within the extended SNX5 PX domain to aspartate prevented the interaction with NRP1, consistent with its inhibitory impact on CI-MPR binding^20^ (**Fig. 4C**). The cytosolic tail of NRP1 contains a stretch of residues (894-918), conserved in NRP1 and NRP2, conforming to the ФxΩxФx_n_Ф motif for SNX5/SNX6 PX domain binding (**Extended Data Fig. 4G**). Isothermal titration calorimetry (ITC) showed that a synthetic peptide corresponding to these residues in NRP1 directly bound the PX domain of SNX5 and SNX6 with an affinity of 19.8 µM and 14.3 µM respectively, which is similar in strength to other known cargoes^20^ (**Fig. 4D** **and Extended Data Fig. 4G**). Next, using structures of ФxΩxФx_n_Ф cargoes bound to the SNX5 PX domain^20^, we generated a molecular model for the NRP1:SNX5 PX interaction. This was consistent with the NRP1 cytosolic tail folding into a beta hairpin that docked into the SNX5 PX domain (βA: 898-NYNFELV-904, loop: 905-DG-906, βB: 907-VKLKKD-912) (**Fig. 4E**).

To validate the model, we generated a panel of mutants of the residues of the predicted βA and βB in the GFP-tagged cytosolic tail of NRP1 (**Extended Data Fig. 4H**). Immunoprecipitation experiments revealed that the mutant forms of the NRP1 tail displayed a reduced interaction with mCherry-SNX6 (**Extended Data Fig. 4I-J**). ITC experiments confirmed a dramatic loss of affinity of the NRP1 tail for the PX domain of SNX5 when the aromatic residues Y899 and F901 were mutated (**Extended Data Fig. 4K**). Furthermore, a triple deletion of the βA residues ^898^NYN^900^ produced the largest reduction in mCherry-SNX6 immunoprecipitation and endogenous SNX5 and SNX6 immunoprecipitation (**Fig. 4F**). Importantly, deletion of these residues did not affect PSD-95/Dlg/ZO-1 (PDZ)-binding motif-mediated association with GIPC1, the scaffolding protein that regulates NRP1 internalisation through binding to a C-terminal PDZ-binding motif in NRP1^29–31^ (**Extended Data Fig. 4L**).

We next compared the trafficking of GFP-Nrp1 and GFP-Nrp1 ΔNYN in previously characterised NRP1 KO HeLa cells^23^. Although the total and surface levels of GFP-Nrp1 and GFP-Nrp1 ΔNYN Nrp1 were comparable, the retrograde trafficking of the mutant was reduced when compared to GFP-Nrp1 (**Fig. 4G**, **Extended Data Fig. 4M**) and resulted in the increased degradation of the mutant receptor (**Fig. 4H**). Consistently, in SNX5+6 double KO cells, GFP-tagged SNX5 rescued NRP1 turnover rate to wild-type levels, whereas rescue with GFP-SNX5(F136D), which does not immunoprecipitate NRP1, failed to do so (**Fig. 4C,I**). These data establish that NRP1 possesses a canonical ESCPE-1 binding motif, and that direct interaction between NRP1 and ESCPE-1 is required for the correct retrograde trafficking of the internalised receptor.

We recently established that the extracellular b1 domain of NRP1 directly associates with a multibasic C-terminal motif (termed the C-end Rule motif^32^) in the furin-processed SARS-CoV-2 Spike S1 subunit to facilitate infectivity^23^. Therefore, we investigated whether ESCPE-1 could associate with the SARS-CoV-2 Spike (S) protein via NRP1. In HEK293T cells stably expressing a C-terminally truncated SARS-CoV-2 S gene (SARS-2 SΔ1256-1273), GFP-SNX5 and GFP-SNX6 co-immunoprecipitated bands corresponding to S1 and uncleaved S alongside endogenous NRP1 (**Fig. 5A,B**). SARS-CoV-2 S does not contain a putative ФxΩxФx_n_Ф motif within its cytosolic tail, suggesting that co-immunoprecipitation likely occurs through an intermediate protein. Importantly, the association between SNX5 and Spike was abrogated by the SNX5(F136D) mutation incapable of binding NRP1, consistent with Spike binding being mediated through an NRP1-dependent mechanism (**Fig. 5B**).

**Fig. 5.**
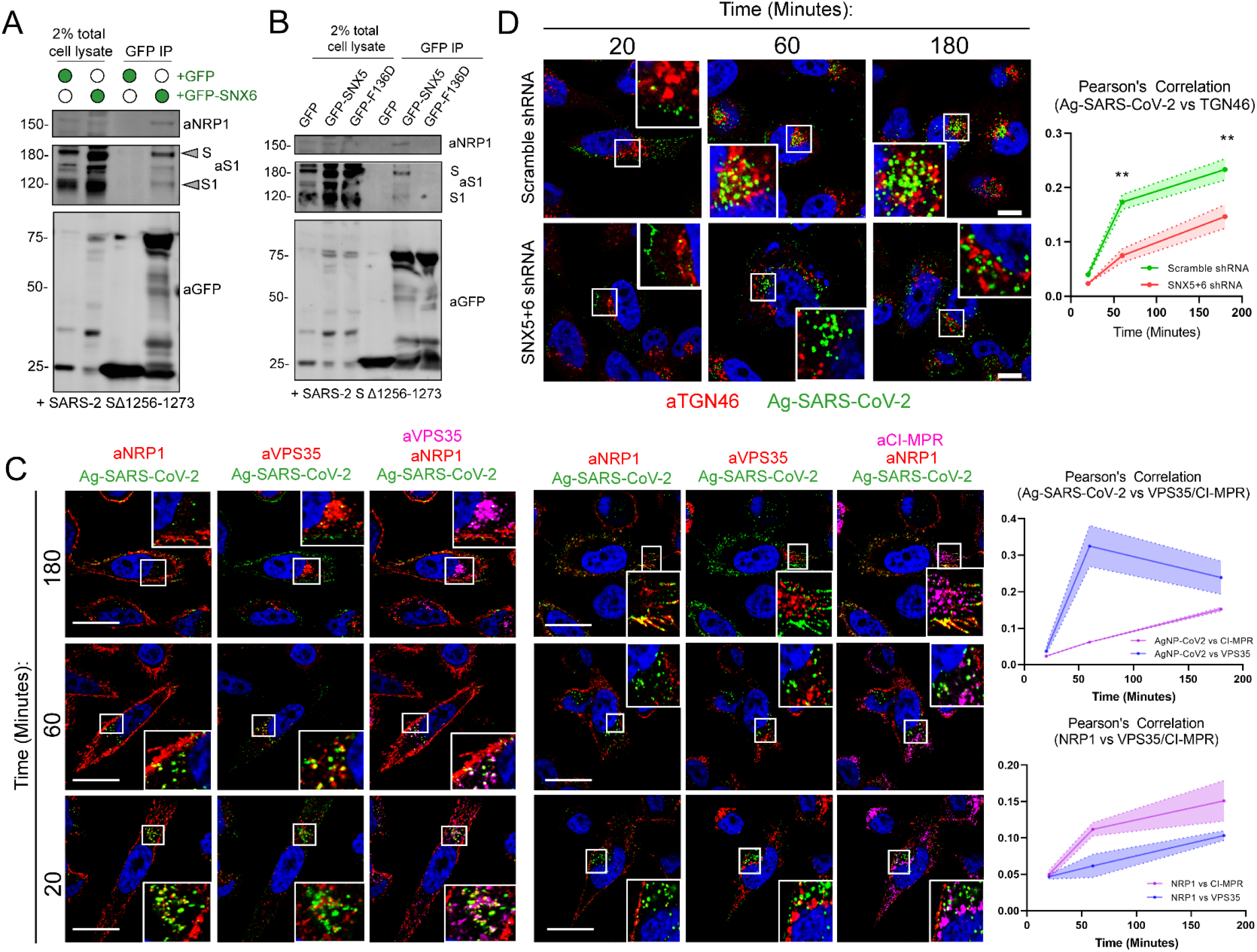
ESCPE-1 Interacts with the SARS-CoV-2 Spike Protein through a Cargo-Selective Mechanism and Mediates Trafficking of Nanoparticles Coated with the SARS-CoV-2 Spike CendR Motif. **(A)** HEK293T cells stably expressing SARS-CoV-2 S (SARS-2 SΔ1256-1273) were transfected to express GFP-tagged SNX6 or GFP and subjected to GFP-nanotrap. **(B)** HEK293T cells stably expressing the SARS-CoV-2 S gene (SARS-2 SΔ1256-1273) were transfected to express GFP-tagged SNX5 or SNX5(F136D) and subjected to GFP-nanotrap. **(C)** Ag-SARS-CoV-2 uptake assay in PPC-1 cells. Pearson’s correlation between Ag-SARS-CoV-2 and anti-VPS35 or anti-CI-MPR is measured over time using Image J software (N = 3 experiments). Scale = 20 µm**. (D)** Ag-SARS-CoV-2 uptake assay in Scramble shRNA PPC-1 cells or SNX5+6 shRNA PPC-1 cells. Pearson’s correlation between Ag-SARS-CoV-2 and anti-TGN46 is measured over time using Image J software (N = 3 experiments). Two-way ANOVA with Šídák’s multiple comparisons test, p = 0.8154 (20 minutes), p = 0.0013 (60 minutes), p = 0.0034 (180 minutes). Scale = 20 µm.

To investigate whether the interaction between NRP1 and ESCPE-1 may regulate retrograde trafficking of internalised NRP1-bound viral particles, we engineered a minimal system comprising approximately coronavirion-sized (80 nm) silver nanoparticles coated with the C-end Rule (CendR) peptide motif of the SARS-CoV-2 S1 protein (Ag-SARS-CoV-2, sequence: TNSPRRAR, original isolate)^23,24^. Silver nanoparticles decorated with CendR sequences specifically bind NRP1 at the cell surface, and are internalised through an NRP1-dependent mechanism^24,32,33^. Accordingly, NRP1-expressing PPC-1 cells incubated with Ag-SARS-CoV-2 demonstrated colocalisation with NRP1 at the cell surface at early timepoints (**Fig. 5C**). Adherence at the cell surface was followed by internalisation of nanoparticles and transit towards the perinuclear region, visualised by early colocalisation with the endosomal marker VPS35 and accumulating colocalisation with CI-MPR and TGN46 over 3 hours (**Fig. 5C,D**). This phenomenon was ESCPE-1 dependent, as SNX5+SNX6 KO PPC-1 cells demonstrated a significant decrease in Ag-SARS-CoV-2 colocalisation with TGN46 and Golgin-97 (**Fig. 5D, Extended Data Fig. 5A**). These data suggest that ESCPE-1 can govern the endosomal trafficking of internalised NRP1 ligands, raising the possibility of a role for the complex in SARS-CoV-2 infection **(Extended Data Fig. 5B**).

## Discussion

We and others recently demonstrated that SARS-CoV-2 directly interacts with NRP1 at the cell surface to enhance cellular infection^23,24^. In the present study, through the development of an unbiased proteomic screen to discover novel retrograde endosomal cargoes, we identified a range of potential transmembrane cargoes for the evolutionarily conserved ESCPE-1 complex. From this screen, we validate NRP1 as an interactor of ESCPE-1. ESCPE-1 directly engages the cytosolic tail of NRP1 in a sequence-dependent manner and mediates tubulovesicular endosomal sorting of NRP1 to the TGN. Additionally, we identify a wider functional network of transmembrane proteins perturbed by SNX5+SNX6 depletion, including integrin-α5 and integrin-β8, and the receptor tyrosine kinases MET and EPHA2, raising the possibility that ESCPE-1-mediated retrograde sorting may play a role in directional cell migration and signalling^34–36^. NRP1 has a reported role in mediating integrin internalisation and trafficking^30,37^, and associates with receptor tyrosine kinases including the MET receptor to modulate their signalling outputs^38,39^. Our study thus provides new mechanistic insight into the previously described role of retrograde endosomal sorting in cell migration^10,36^.

Considering the recent identification of NRP1 as a SARS-CoV-2 host factor, the mechanistic basis of its intracellular trafficking by ESCPE-1 opens interesting avenues for future investigation. SARS-CoV-2 exhibits two distinct cellular entry pathways: either direct fusion with the plasma membrane or internalisation into endosomal compartments^40^. It is presently unknown whether the SARS-CoV-2 virions that are internalised through endocytosis can hijack NRP1- and ESCPE-1-dependent endocytic trafficking to subvert innate cellular defenses, or whether the Spike protein alone undergoes this trafficking step. We demonstrate that ESCPE-1 co-immunoprecipitates the SARS-CoV-2 Spike through a sequence-dependent mechanism likely dependent on NRP1. Furthermore, engineered nanoparticles displaying the CendR motif of SARS-CoV-2 Spike hijacked ESCPE-1-dependent retrograde trafficking. ESCPE-1 couples to the dynein-dynactin complex to provide a mechanical pulling force that aids the biogenesis of tubular endosomal carriers that traffic towards the perinuclear region^25–27^. This process may conceivably provide additional energy input and membrane tension to facilitate the virus endosomal uncoating, as seen in influenza A virus^41,42^. Indeed, modelling of the SARS-CoV-2 Spike protein binding to NRP1 and ACE2 has suggested that NRP1 facilitates S1/S2 separation, a prerequisite for membrane fusion^43^.

Interestingly, additional pathogens hijack ESCPE-1 cargo recognition to promote intracellular survival^2,3,44^, and SNX5 has also recently been identified as a key regulator of innate cellular immunity against a range of viruses^45^. Genome wide CRISPR screens and biochemical studies have identified multiple endosomal sorting machineries that facilitate SARS-CoV-2 infection, including components of the retromer, retriever and COMMD/CCDC22/CCDC93 (CCC) complexes^46–51^. Here, we suggest that ESCPE-1 sequence-dependent cargo sorting also plays a role in regulating the endosomal dynamics exploited during SARS-CoV-2 infection, potentially expanding the scope of pathogens that exploit can this complex.

A fascinating and impactful emerging theme is that the NRP1 pathway may influence infection by a wide range of viruses through the recognition of CendR motifs on viral glycoproteins^32,52^. Our findings therefore highlight the possibility that multiple viruses converge upon a NRP1- and ESCPE-1-dependent intracellular trafficking pathway within the endosomal network. We previously demonstrated that pharmacological inhibition of the SARS-CoV-2-NRP1 interaction limits infection in cell culture^23^. Future work will be required to appreciate the importance of NRP1 trafficking in SARS-CoV-2 biology, and the wider range of pathogens that exploit this receptor to mediate infectivity.

## Acknowledgements

We thank Guido Serini (Candiolo Cancer Institute - Fondazione del Piemonte per l’Oncologia (FPO) - IRCCS, Candiolo (TO), Italy) and Donatella Valdembri (Department of Oncology, University of Torino School of Medicine, Candiolo (TO), Italy) for their gift of the GFP-tagged Nrp1 construct and invaluable discussion. We thank the Wolfson Bioimaging Facility and the Proteomics Facility at the University of Bristol for their support. We thank the University of Bristol Advanced Computing Research Centre for the provision of high performance computing.

P.J.C is supported by the Wellcome Trust (104568/Z/14/Z and 220260/Z/20/Z), the Medical Research Council (MR/L007363/1 and MR/P018807/1), the Lister Institute of Preventive Medicine, and the award of a Royal Society Noreen Murray Research Professorship to P.J.C. (RSRP/R1/211004). J.L.D was supported by a Wellcome Trust studentship from the Dynamic Molecular Cell Biology Ph.D program (203959/Z/16/Z), C.A.P was supported by Beca Fundación Ramón Areces Estudios Postdoctorales en el Extranjero. D.K.S was supported by a BrisSynBio BBSRC research grant (BB/L01386X/1). B.M.C. is supported by a Senior Research Fellowship and Project Grant from the National Health and Medical Research Council (APP1136021 and APP1156493). Y.Y. is supported by the European Research Council under the European Union’s Horizon 2020 research and innovation program (No 856581 – CHUbVi). A.D.D is supported by the UK Research and Innovation/Medical Research Council (MRC) (MR/W005611/1). TT was supported by the European Regional Development Fund (Project No. 2014-2020.4.01.15-0012), by Estonian Research Council (grant PRG230 and EAG79) and ERA-NET EuroNanoMed III project iNanogun.

## Author Contributions

Initial concept: B.S, J.L.D and P.J.C. Experimental design and analysis: B.S, J.L.D, L.S.G, K.K, S.W, C.A.P, A.T, L.H, P.L, K.J.H, D.K.S. Supervision and funding acquisition: B.S, A.D.D, B.M.C, T.T, Y.Y and P.J.C. Manuscript writing: B.S, J.L.D and P.J.C. Manuscript review and editing: all authors.

## Material and Methods

### Antibodies

Antibodies used in this study were: GFP (clones 7.1 and 13.1; 11814460001; Roche) (1:2000 for WB, 1:400 for IF), SNX1 (clone 51/SNX1; 611482; BD) (1:1000 for WB, 1:200 for IF), SNX2 (clone 13/SNX2; 5345661; BD) (1:1000 for WB, 1:200 for IF), SNX6 (clone d-5,365965; Santa Cruz Biotechnology, Inc.) (1:1000 for WB, 1:200 for IF), SNX5 (clone EPR14358; ab180520; Abcam) (WB 1:500) , β-actin (A1978; Sigma-Aldrich) (1:2000 for WB), ITGA5 (610633, BD) (1:1000 for WB); VPS35 (97545; Abcam) (IF 1:200), GFP (GTX20290; GeneTex) (WB 1:2000), EEA1 (C45B10; Cell Signalling Technologies) (IF 1:200), LAMP1 (clone H4A3, AB2296838; DSHB) (IF 1:200) Cathepsin D (21327-1-AP; Proteintech) (WB 1:1000), Calnexin (ab22595; Abcam) (WB 1:1000), TGN46 (AHP500G, Bio-Rad) (IF 1:400), Golgin-97 (Clone CDF4; A-21270; Thermo Fischer) (IF 1:200), HA tag (66006-1-Ig, Proteintech) (WB 1:1000), NCAD (Clone 13A0; 14215; Cell Signalling) (WB 1:1000), Tom20 (Clone 29; 612278; BD Biosciences), GALNT2 (ab102650; Abcam) (WB 1:1000), NRP1 (Clone EPR3113; ab81321; Abcam) (WB 1:1000, IF 1:200), NRP1 b1b2 domain^23,24^ (Clone 3E7.1) (IF 1:200), Tubulin (ab6046; Abcam) (WB 1:2000). SARS-CoV-2 N (200-401-A50, Rockland) (IF 1:2000), SARS-CoV-2 Spike (40592-T62, Sino Biologicals) (WB 1:1000).

### Cell culture and Transfection

HeLa and HEK-293T cell lines were sourced from ATCC. Authentication was from the ATCC. PPC-1 human primary prostate cancer cells were obtained from Erkki Ruoslahti laboratory at Cancer Research Center, Sanford Burnham-Prebys Medical Discovery Institute. Cells were grown in DMEM (Sigma-Aldrich) supplemented with 10% (v/v) FCS (Sigma-Aldrich) and penicillin/streptomycin (Gibco) and grown under standard conditions. FuGENE HD (Promega) was used for transient transfection of DNA according to the manufacturer’s instructions. Cycloheximide was used to block protein synthesis. Cycloheximide (C7698; Sigma-Aldrich) at 10 µg/ml was added to the cells for the indicated time points. The SNX1+SNX2 and SNX5+SNX6 knock-out clonal cell lines, and HeLa^wt^ +ACE2 and HeLa^NRP1 KO^ + ACE2 cell lines used in this study were characterised previously^19,20,23^. For GFP-based immunoprecipitations, HEK293T cells were transfected with GFP constructs using polyethylenimine (Sigma-Aldrich) and expression was allowed for 48 hours. For siRNA-based knockdown, cells were first reverse-transfected using DharmaFECT (GE Healthcare) and then transfected again with HiPerFect (QIAGEN) 24 hours later according to the manufacturer’s instructions. 48 hours after the second transfection, cells were lysed or fixed and stained. SNX5+6 suppression was performed using a combination of the following oligonucleotides against SNX5 (sequence 5′-CUACGAAGCCCGACUUUGA-3′) and SNX6 (sequence 5′-UAAAUCAGCAGAUGGAGUA-3′). To perform knockdowns in PPC-1 cells, pLKO.1-puro-CMV-tGFP plasmids targeting SNX5 (5’-ACTATTACAATAGGATCAAAG-3’, 5’-CTGAGTATCTCGCTGTGTTTA-3’) and SNX6 (5’-AGTAAAGGATGTAGATGATTT-3’, 5’-GCCGAAACTTCCCAACAATTA-3’) (Sigma-Aldrich) were lentivirally transduced, and GFP-positive cells quantified as knockdowns. HeLa cells were transduced with lentiviruses, with constructs cloned into pXLG3 or pLVX vectors,to produce stably expressing cell lines. Following transduction, transduced cells were selected with puromycin or blasticidin accordingly.

### Immunoprecipitation and Quantitative Western Blot Analysis

For western blotting, cells were lysed in PBS with 1% (v/v) Triton X-100 and protease inhibitor cocktail. The protein concentration was determined with a BCA assay kit (Thermo Fisher Scientific), and equal amounts were resolved on NuPAGE 4–12% precast gels (Invitrogen). Blotting was performed onto polyvinylidene fluoride membranes (Immobilon-FL; EMD Millipore) followed by detection using the Odyssey infrared scanning system (LI-COR Biosciences). For GFP-based immunoprecipitations, HEK-293T cells were lysed 48 h after transfection in immunoprecipitation buffer (50 mM Tris-HCl, 0.5% (v/v) NP-40, and Roche protease inhibitor cocktail) and subjected to GFP trap (ChromoTek). Immunoblotting was performed using standard procedures. Detection was performed on an Odyssey infrared scanning system (LI-COR Biosciences) using fluorescently labeled secondary antibodies.

### HRP-TGN46 Proximity Biotinylation

The methodology for HRP-TGN46 biotinylation is adapted from the APEX2 biotinylation protocol outlined in^15^. 10×10^6^ HRP-TGN46-expressing cells were seeded in a 15 cm plate the day before biotinylation. The next day, cells were incubated in DMEM media supplemented with 500 μM biotin-phenol (BP) and incubated for 30 minutes at 37°C. Hydrogen peroxide (H_2_O_2_) was added at a final concentration of 1 mM and distributed by rocking the cell plate. After 45 seconds of H_2_O_2_ incubation, the media was removed and replaced with ice-cold, freshly prepared quencher solution consisting of 1 mM sodium ascorbate, 500 μM (±)-6-Hydroxy-2,5,7,8-tetramethylchromane-2-carboxylic acid (Trolox, Sigma-Aldrich, 238813) and 1 mM sodium azide in PBS to ensure that the biotinylation reaction does not proceed beyond 1 minute. The quencher solution was left for 1 minute, then discarded, and this washing process was repeated 5 times. Following washes, cells were lysed in RIPA buffer (150 mM NaCl, 0.1% (v/v) SDS, 0.5% (w/v) sodium deoxycholate, 1% (v/v) Triton X-100, protease inhibitor cocktail, 50 mM Tris-HCl, pH 7.5) and lysates spun at 20,000 x g for 10 minutes at 4°C. 50μL aliquots of streptavidin beads were prepared and washed 3 times in RIPA buffer, centrifuging beads between washes at 350 x g. The cell lysates were mixed with the streptavidin beads and rotated for 1 hour at 4°C.

After incubation, streptavidin beads were centrifuged and the supernatant containing unbound proteins is removed. The beads were washed 7 times (twice in RIPA buffer, once with 1M KCl, once with 0.1M Na_2_CO_3_, once with 2M Urea 10mM Tris-HCl pH 8.0, and twice again with RIPA buffer). All buffers were kept ice cold throughout the process. After the final wash step, all supernatant was aspirated off and beads were resuspended in 3X NuPAGE sample buffer supplemented with 2.5% β-mercaptoethanol, 2mM free biotin and 20mM DTT.

### Biotinylation of Cell Surface Proteins

For surface biotinylation experiments, fresh Sulfo-NHS-SS Biotin (Thermo Scientifics, #21217) was dissolved in ice-cold PBS at pH 7.8 at a final concentration of 0.2 mg / ml. Cells were washed twice in ice-cold PBS and placed on ice to slow down the endocytic pathway. Next, cells were incubated with the biotinylation reagent for 30 minutes at 4°C followed by incubation in TBS for 10 minutes to quench the unbound biotin. The cells were then lysed in lysis buffer and subjected to Streptavidin beads-based affinity isolation (GE-Healthcare).

### Immunofluorescence Staining

Cells were fixed in 4% (v/v) PFA for 20 min and washed three times in PBS and permeabilised with 0.1% (v/v) Triton X-100. Fixed cells were blocked in 1% (w/v) BSA and incubated in primary antibody and respective secondary antibody (Alexa Fluor; Thermo Fisher Scientific) in 1% (w/v) BSA. Biotinylated proteins were labelled with 0.5 μg/mL Alexa Fluor 568-conjugated streptavidin. For uptake assays, HeLa cells were transfected with GFP-Nrp1 constructs. 24 h after transfection, cells were incubated with anti-GFP antibody on ice for 30 minutes, then returned to 37°C media. GFP-Nrp1 trafficking was followed for 0 hours, 1 hours, 4 hours and 8 hours prior to fixation. The retrograde transport to the TGN was assayed through measuring colocalisation of anti-GFP signal with the TGN marker TGN46.

### Electron Microscopy

100,000 HRP-TGN46-expressing HeLa cells were seeded in glass bottom 35 mm dishes (MatTek, P35G-1.5-14-CGRD) the day before sample processing. Cells were fixed in solution of 2.5% glutaraldehyde, 3 mM CaCl_2_, 0.1 M cacodylate buffer, pH 7.4, for 5 minutes at room temperature, then 1 hour on ice. All solutions and incubation steps from this point onwards until the embedding stage were kept ice cold. Samples were washed 5 times in 0.1 M cacodylate buffer, leaving each wash on ice for 2 minutes. Samples were incubated in a quenching buffer of 20 mM glycine and 0.1 M cacodylate for 5 minutes, then washed in 0.1 M cacodylate buffer a further 5 times, at 2 minutes per wash. A 1X diaminobenzidine (DAB), 10 mM H_2_O_2_ solution was assembled by dissolving 50 mg of DAB in 10 mL of HCl, then diluting 1 mL of this solution in 9 mL of 0.1 M cacodylate and 10 µL of 30% (w/w) H_2_O_2_.

To obtain differential interference contrast (DIC) images of DAB polymerisation, a Leica DM IRBE inverted epifluorescence microscope (Leica Microsystems) was used. An initial picture was taken prior to DAB labelling. The 0.1 M cacodylate washing buffer was removed and replaced with the 1mL of 1X DAB 10 mM H_2_O_2_ solution. DAB polymerisation was observed through the eyepiece in real time, and DIC images were taken. The reaction was stopped by removing the solution and washing 5 times with 0.1 M cacodylate buffer, at 2 minutes per wash. The final wash was removed, and cells were incubated in 1% OsO_4_ for 30 minutes. This solution was removed, then samples were washed with distilled water 3 times, at 1 minute per wash. Finally, samples were incubated overnight in a filtered solution of 2% uranyl acetate in distilled water at 4°C.

The next day, samples were dehydrated by sequential 3-minute incubations of 20%, 50%, 70%, 90%, 100%, 100% ice cold ethanol, followed by a final 3 minute wash of 100% ethanol at room temperature. The final ethanol wash was removed and EPON resin (TAAB, T031) was poured onto the samples, then samples were left rocking for 3 hours. The resin was discarded, then fresh EPON resin was poured onto samples. The samples were incubated at 60°C overnight to set the resin. A small volume of fresh EPON was poured into the middle of the sample, then used to adhere an EPON stub. The sample was returned to 60°C for a further 24 hours. The coverslip was removed from the resin by sequential plunging into liquid nitrogen and boiling water. The embedded resin was cut into < 70 nm slices for transmission electron microscopy using a Leica UC6 cryo ultra microtome (Leica Microsystems). Samples were imaged on a FEI Tecnai 12 120 kV BioTwin Spirit transmission electron microscope (FEI Company).

### Image Acquisition and Image Analysis

Microscopy images were collected with a confocal laser-scanning microscope (SP5 AOBS; Leica Microsystems) attached to an inverted epifluorescence microscope (DMI6000; Thermo Fisher Scientific). A 63× 1.4 NA oil immersion objective (Plan Apochromat BL; Leica Biosystems) and the standard SP5 system acquisition software and detector were used. For stimulated emission depletion (STED) microscopy, images were taken using a Leica SP8 confocal laser scanning microscope attached to a Leica DMi8 inverted epifluorescence microscope (Leica Microsystems) with a 63x HC PL APO CS2 oil immersion lens, numerical aperture 1.4 (Leica Microsystems, 506351). Images were captured at room temperature as z stacks with photomultiplier tube detectors with a photocathode made of gallium-arsenide-phosphide (Leica Microsystems) for collecting light emission. Images were captured using Application Suite AF software (version 2.7.3.9723; Leica Microsystems) and then analyzed with the Volocity 6.3 software (PerkinElmer) or ImageJ. For live cell imaging, cells were seeded in dishes (MatTek) in prewarmed CO2-independent media. Cells were imaged with a confocal laser scanning microscope (SP8 AOBS; Leica Microsystems) attached to a DMI6000 inverted epifluorescence microscope with an HCX Plan Apochromat lambda blue 63× 1.4 NA oil objective. Images were captured at 37°C, and “Adaptive Focus Control” was used to correct focus drift during time courses.

### Statistics and Reproducibility

All quantified Western blot are the mean of at least three independent experiments. Statistical analyses were performed using GraphPad Prism 9 (LaJolla, CA). Graphs represent means and S.E.M. For all statistical tests, p < 0.05 was considered significant and is indicated by asterisks.

### Stable Isotope Labelling of Amino Acids in Cell Culture (SILAC)-Based Proteomics

HeLa cells expressing HRP-TGN46 were cultured for at least 6 doublings in three different isotopically labelled media compositions: R_0_K_0_ (light), R_6_K_4_ (medium), R_10_K_8_ (heavy) (Silantes). After performing HRP-TGN46 biotinylation and streptavidin affinity purification, the streptavidin beads corresponding to different SILAC conditions were pooled together prior to washing steps. Biotinylated proteins were eluted in a volume of 40 μL 3X NuPAGE sample buffer supplemented with 2.5% β-mercaptoethanol, 2mM free biotin and 20mM DTT. The eluate was loaded onto a gel, and proteins were resolved by SDS-PAGE, then visualised by staining with SimplyBlue SafeStain (Thermo Fisher, LC6060). The gel lane was cut into 10 equal slices, which were subjected to in-gel tryptic digestion and the resulting peptides analysed by nano-liquid chromatography tandem mass spectrometry (LC-MS/MS) using an Orbitrap Fusion Tribrid mass spectrometer (Thermo Fisher Scientific).

The raw data files were processed and quantified using Proteome Discoverer software v2.1 (Thermo Fisher Scientific) and searched against the UniProt Human database using the SEQUEST HT algorithm. Peptide precursor mass tolerance was set at 10ppm, and MS/MS tolerance was set at 0.6Da. Search criteria included carbamidomethylation of cysteine as a fixed modification and oxidation of methionine, appropriate SILAC labels and the addition of biotin-phenol (+361.146Da) to tyrosine as variable modifications. Searches were performed with full tryptic digestion and a maximum of 4 missed cleavages were allowed. The reverse database search option was enabled and all data was filtered to satisfy false discovery rate (FDR) of 5%.

For statistical analysis of differential protein abundance between conditions, standard t-tests were used. Volcano plots were plotted using Orange software (University of Ljubljana) or VolcanoseR^53^. For generation of the HRP-TGN46-labelled proteome, proteins that were only identified in the HRP-TGN46 biotinylation condition in ≥ 4 out of 5 repeats and were not identified in negative control conditions and thus could not be statistically analysed, were assumed to be significant hits (**Supplementary Table 2**). Furthermore, 10 proteins significantly enriched in the heavy SILAC condition (HRP-TGN46 + BP – H_2_O_2_) relative to the light SILAC condition (WT HeLa + BP + H_2_O_2_) were identified and removed from downstream analyses.

Gene ontology analysis was performed using the PANTHER classification system^54^ and Metascape^55^. Protein IDs for enriched or depleted proteins were compared against the total human genome. Gene ontology terms, falling under the categories of ‘Cellular Component’, ‘Biological Process’ or ‘Molecular Function’ that were significantly enriched or depleted relative to the expected number for the sample size of proteins were identified by a Fisher’s exact test with the Bonferroni correction for multiple testing by PANTHER software.

### Recombinant Protein Expression and Purification

The expression plasmid of pGEX-4T-2 containing GST tagged SNX5 and SNX6 PX domain constructs were transformed into Escherichia coli BL 21 (DE3) cells and plated on lysogeny-broth (LB) agar plates supplemented with Ampicillin (0.1mg/mL). Single colony was then used to inoculate 50 mL of LB medium containing Ampicillin and the culture was grown overnight at 37 °C with shaking at 180 rpm. The following day, 1L of LB medium containing antibiotics Ampicillin (0.1mg/mL) was inoculated using 10 ml of the overnight culture. Cells were then grown at 37 °C with shaking at 180 rpm to an optical density of 0.8-0.9 at 600 nm and the protein expression was induced by adding 0.5 mM IPTG (isopropyl-b-D-thiogalactopyranoside). Expression cultures were incubated at 20 °C overnight and the cells were harvested next day by centrifugation at 4000 rpm for 15 min using Beckman rotor JLA 8.100. Cell pellets were then resuspended in 20 mL (for cell pellet from 1L) of lysis buffer (50 mM HEPES, pH 7.5, 500 mM NaCl, 5% glycerol, Benzamidine (0.1mg/mL), and Dnase (0.1 mg/mL)). Resuspended cells were lysed by using the cell disrupter (Constant systems, LTD, UK, TS-Series) and the soluble fraction containing the protein was separated from cell debris by centrifugation at 18,000 rpm for 30 min at 4 °C. The soluble fraction was first purified by affinity chromatography using Glutathione Sepharose 4B resin (GE Healthcare) and the GST tag was cleaved by incubating the protein with Thrombin (Sigma Aldrich) overnight at 4 °C. Next day the protein was eluted using 50 mM HEPES, pH 7.5, 200 mM NaCl. The eluted protein was then concentrated and further purified by gel filtration chromatography (Superdex 75 (16/600), GE Healthcare) using 50 mM HEPES, pH 7.5, 200 mM NaCl, 0.5 mM TCEP (tri(2-carboxyethyl)phosphine) and the fractions corresponding to SNX5/SNX6 PX were analysed by SDS PAGE.

### Isothermal Titration Calorimetry (ITC)

ITC experiments were carried out by using Microcal ITC200 instrument (Malvern) at 298 K. The NRP1 peptide was dissolved in the same buffer (50 mM HEPES, pH 7.5, 200 mM NaCl, 0.5 mM TCEP) as the proteins to avoid buffer mismatch. 25 mM SNX5/6 PX was then titrated with 1 mM NRP1 peptide using a series of 13 injections of 3.22 ml each with 180 s intervals. The dissociation constant, *K*_d_ (1/*K*_a_), enthalpy of binding (Δ*H*)) and the stoichiometry of the binding reaction (N) were calculated by fitting the data to a single site binding model using MicroCal PEAQ-ITC software. Experiments were performed in triplicate to check for reproducibility and the average values of the three experiments are reported.

### SNX5-NRP1 Modelling and Molecular Dynamics

The model for the SNX5-NRP1 complex was built based on the 6n5z.pdb structure according to the method outlined in Supplementary Information. 20 ns atomistic dynamic simulations of the modelled complexes were carried out using the amber99sb-ildn forcefield in TIP3P waters and GROMACS^56^ (2019.2) according to the method described recently^57^.

### Silver Nanoparticles Uptake Assay

Silver nanoparticles labelled with the dye CF555 and functionalised with the biotinylated CendR peptide of the SARS-CoV-2 S1 protein (biotin-Ahx-TNSPRRAR; Ahx = aminohexanoic acid) were prepared as previously described^58,59^. The peptide was purchased from TAG Copenhagen, Copenhagen, Denmark.

PPC-1 cells (10^5^ cells) were seeded onto noncoated coverslips (12 mm diameter, Marien-feld-Superior, Paul Marienfeld GmbH & Co.KG, Lauda-Königshofen, Germany) in a 24-well plate and cultured for 24 h. The cells were incubated with the mouse monoclonal anti-human NRP1 antibody (clone 3E7.1) at 4 °C for 30 min, washed with cold medium and, subsequently incubated with the AgNPs (0.3 nM in DMEM with 10% FBS) at 37 °C. Cells were washed, treated with the etching solution to remove the non-internalised AgNPs (1 mM of K_3_Fe(CN)_6_ and Na_2_S_2_O_3_ in PBS) at room temperature for 5 min and washed with PBS. For immunosfluorescence staining, cells were fixed in 4% (v/v) PFA for 10 min, washed three times in PBS and permeabilised with 0.2% (v/v) Triton X-100. Fixed cells were blocked in blocking buffer containing 5% BSA (w/v), 5% FBS (v/v), and 5% goat serum (v/v) in PBS with 0.05% Tween-20 (PBST) and incubated in primary antibody and respective secondary antibody (Alexa Fluor; Thermo Fisher Scientific) in blocking buffer diluted 1:5 with PBST. Cells were counterstained with DAPI and visualised using a confocal microscope FV1200MPE (Olympus, Shinjuku, Japan) equipped with the UPlanSApo 60x/1.35na objective (Olympus, Shinjuku, Japan). The images were analysed using Olympus FluoView Ver.4.2a Viewer software. The colocalisation of the AgNPs with the different markers was quantified using the ImageJ software.

---

**Extended Data Fig. 1.**
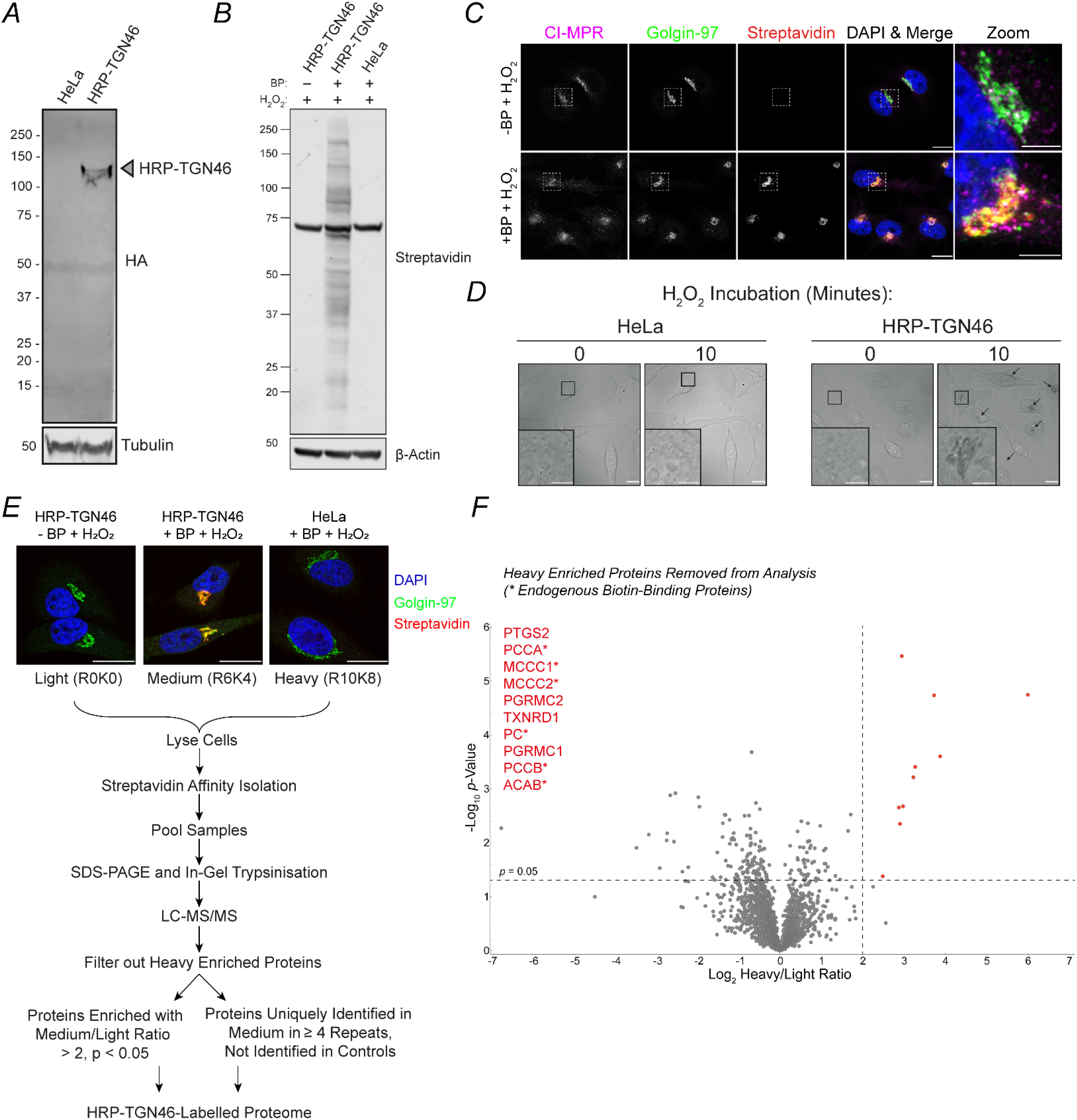
Validation of the HRP-TGN46 Labelling Methodology. **(A)** Identification of HRP-TGN46 in concentrated whole cell lysates with an anti-HA antibody. **(B)** Streptavidin labelling of biotinylated proteins in HRP-TGN46-expressing HeLa cells. **(C)** Fluorescent streptavidin labelling of HRP-TGN46-expressing HeLa cells incubated with H_2_O_2_ in the absence or presence of biotin-phenol. The TGN marker Golgin-97 and retrograde endosomal cargo CI-MPR are labelled by immunofluorescence. Scale bars = 20 µm, 5 µm insets. **(D)** Differential interference contrast imaging of HRP-TGN46 expressing cells before and after incubation with DAB and H_2_O_2_. Arrows indicate electron-dense contrast after 10 minutes localising to the Golgi/TGN. Scale bar: Scale bar: 20 µm, zoom scale bar: 5 µm. **(E)** SILAC experimental design and workflow for the identification of the HRP-TGN46-labelled proteome. Scale bars = 20 µm. **(F)** Volcano plot of proteins identified following streptavidin affinity isolation displayed as a ratio of heavy (HeLa + BP + H_2_O_2_) over light (HRP-TGN46 – BP + H_2_O_2_) abundance. Proteins significantly enriched in the heavy condition (p < 0.05, Log_2_ fold change >2) were filtered out of subsequent analysis. Asterisks represent significantly enriched proteins with known endogenous biotin-binding affinity.

**Extended Data Fig. 2.**
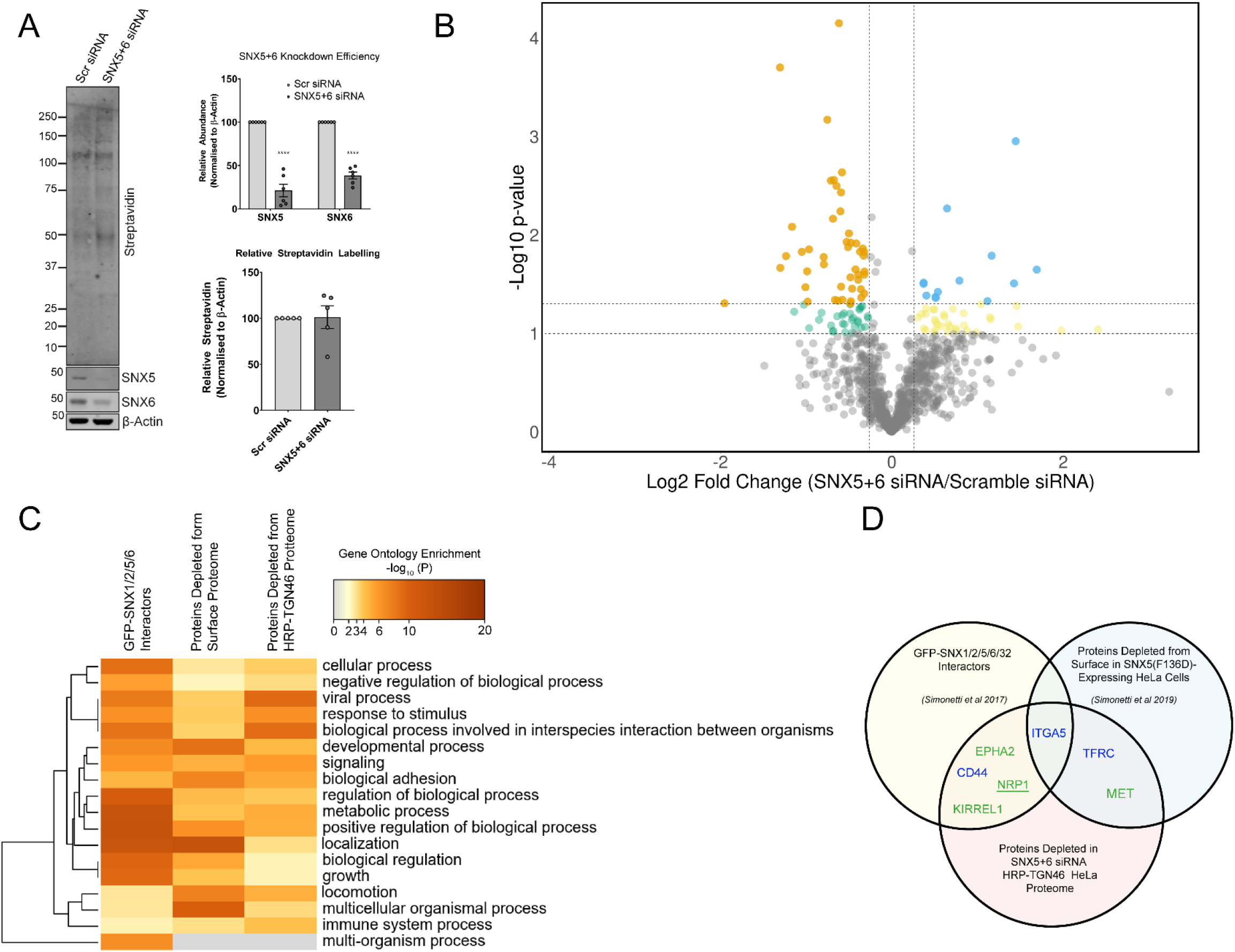
Proteomic Identification of Potential ESCPE-1 Retrograde Cargoes. **(A)** Western blot of whole cell lysates from HRP-TGN46-expressing HeLa cells treated with Scr siRNA or SNX5+6 siRNA. SNX5 and SNX6 levels were quantified following normalisation to β-Actin. 2-way ANOVA with Šídák’s multiple comparisons test, n=6, p < 0.0001 (SNX5), p < 0.0001 (SNX6). Total streptavidin was quantified following normalisation to β-Actin, unpaired t-test, n=5, p = 0.9272. **(B)** Volcano plot displaying HRP-TGN46 target proteins and their fold change as a ratio of SNX5+6 siRNA abundance/Scr siRNA abundance. 76 proteins are highlighted as passing a p < 0.1,Log_2_ fold change < -0.26 threshold (green), and 46 passing a p < 0.05, Log_2_ fold change < -0.26 threshold (orange). **(C-D)** Heat map of gene ontology term enrichment (C) and Venn diagram of overlapping proteins (D) in lists of ESCPE-1 interactors^19^, cell surface cargoes^20^ and proteins depleted in the HRP-TGN46 proteome (p < 0.1, Log_2_ Fold change < -0.26) upon SNX5+6 siRNA suppression.

**Extended Data Fig. 3.**
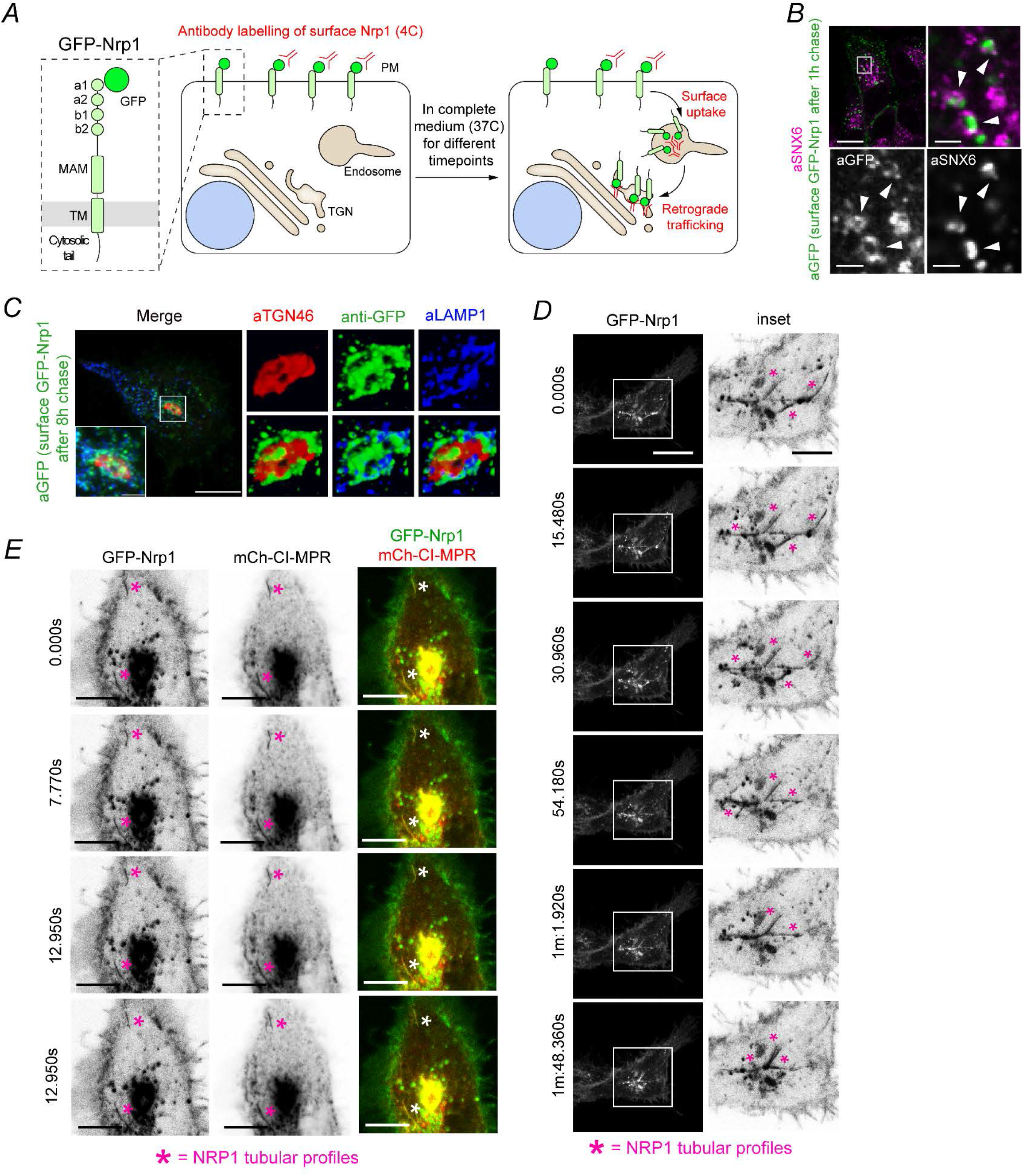
Internalisation and Tubulovesicular Trafficking of a GFP-Nrp1 Construct. **(A)** Schematic of the GFP-Nrp1 uptake assay with the two-step staining procedure used to distinguish total from internalised GFP-Nrp1. **(B)** HeLa cells were transfected with GFP-Nrp1 for 24 hours and subjected to surface labelling of GFP-Nrp1. The uptake assay was followed for 1 hour. Cells were fixed and immunostained for the ESCPE-1 subunit SNX6. Arrowheads: examples of colocalisation of GFP-Nrp1 and SNX6 on tubular endosomes. Scale bar = 20 µm, Inset = 5 µm. **(C)** 8 hour surface uptake of GFP-Nrp1 in HeLa cells. The volume of the perinuclear region, including the TGN was reconstructed from serial z-stacks. Scale bar = 20 µm, Inset = 5 µm. **(D)** HeLa cells were transfected with GFP-Nrp1 and live imaged after 24 hours. Asterisk = examples of Nrp1-positive tubular profiles that emanate from intracellular compartments. Scale bar = 20 µm, Inset = 10 µm. **(E)** HeLa cells were cotransfected with GFP-Nrp1 and mCherry-CI-MPR and live imaged after 24 hours. Asterisk = examples of Nrp1 and CI-MPR signals colocalising in tubular profiles that dynamically move within the cells. Scale bar = 10 µm.

**Extended Data Fig. 4.**
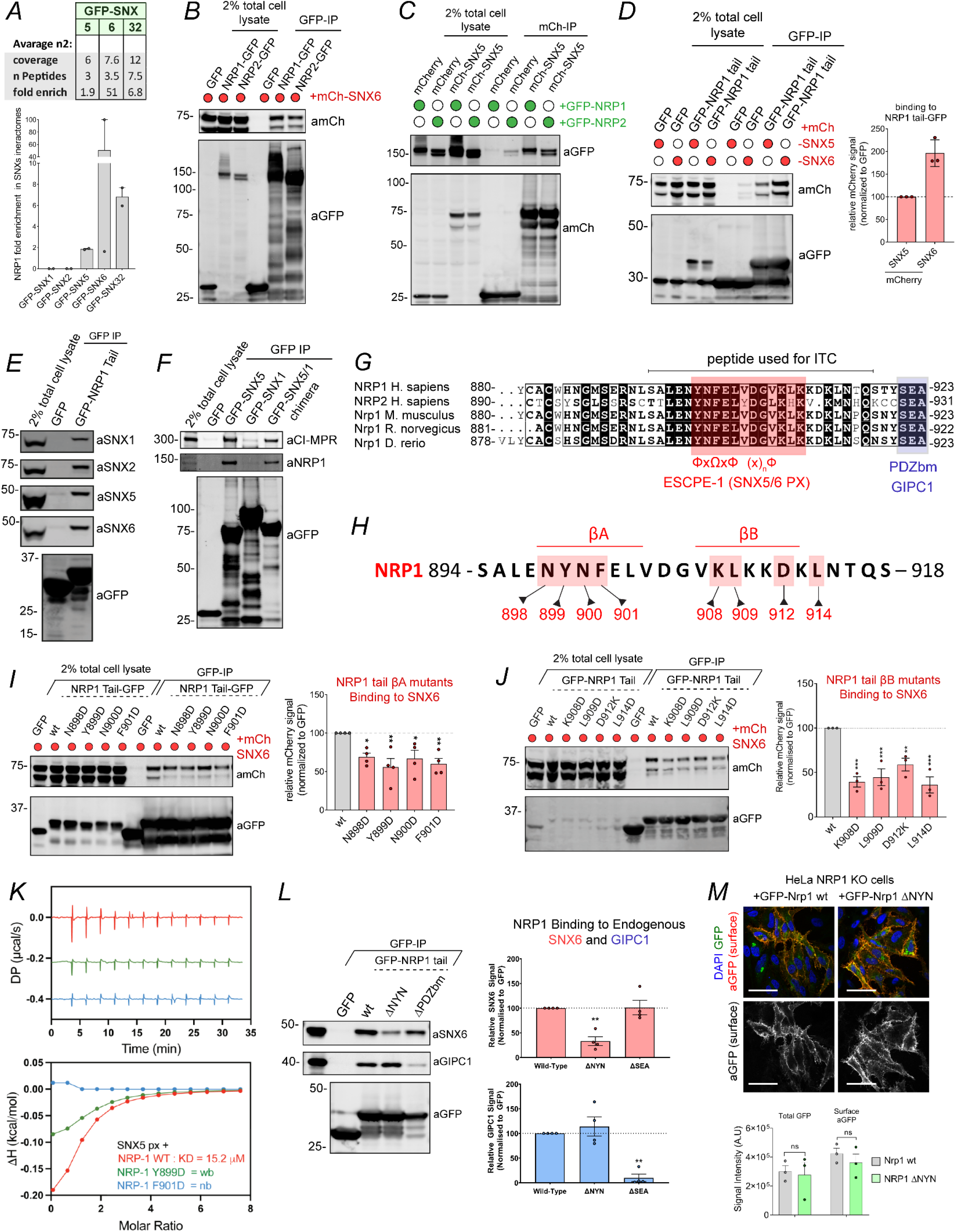
NRP1 and NRP2 Interact with ESCPE-1 Subunits. **(A)** Enrichment and coverage of NRP1 in the previously published SILAC-based proteomics of SNX5, SNX6 and SNX32^19^. **(B)** HEK293T cells were cotransfected to express GFP-tagged NRP1 or NRP2 and mCherry-tagged SNX6 and subjected to GFP-nanotrap. The blot is representative of three independent GFP traps. **(C)** HEK293T cells were cotransfected to express GFP-tagged NRP1 or NRP2 and mCherry or mCherry-tagged SNX5 and subjected to mCherry-nanotrap. The blot is representative of three independent mCherry traps. **(D)** HEK293T cells were cotransfected to express GFP, NRP1-GFP or NRP2-GFP, and mCherry, mCherry-SNX5 or mCherry-SNX6 and subjected to GFP-nanotrap. The band intensity of mCherry was measured from N = 4 independent experiments using Odyssey software. **(E)** HEK293T cells were transfected with GFP or GFP-NRP1 Tail, and lysates subjected to GFP nanotrap and blotting for ESCPE-1 subunits. Data representative of three independent repeats. **(F)** HEK293T cells were transfected to express GFP-tagged SNX5, SNX1 and a SNX1 chimera generated by the replacement of the SNX1 Px domain with that of SNX5. Data representative of three independent repeats. **(G)** Sequence alignment of NRP1 and NRP2 from *H. sapiens*, and Nrp1 from *M. musculus*, *R. norvegicus* and *D. rerio***. (H)** Schematics of the NRP1 cytosolic tail sequence that accounts for the ESCPE-1 binding motif, the residues mutated in the constructs used in this study are highlighted in red. **(I)** HEK293T cells were cotransfected to express GFP-tagged wildtype or mutant forms of the NRP1 tail together with mCherry-SNX6 and subjected to GFP-nanotrap. The band intensity of mCherry was measured from N = 4 independent experiments using Odyssey software. The relative binding of mutant forms over the wild-type NRP1 tail was measured using one-way ANOVA and Dunnett’s test. N898D: p = 0.0454; Y899D: p = 0.0046; N900D: p = 0.0333; F901D: p = 0.0094. **(J)** HEK293T cells were cotransfected to express GFP-tagged wildtype or mutant forms of the NRP1 tail together with mCherry-SNX6 and subjected to GFP-nanotrap. The band intensity of mCherry was measured from N = 3 independent experiments using Odyssey software. The relative binding of mutants forms the wild-type NRP1 tail was measured using one-way ANOVA and Dunnett’s test. K908D: p = 0.0005; L909D: p = 0.0010; D912K: p = 0.0077; L914D: p = 0.0003. **(K)** SNX5 PX domain was titrated against different wild-type, Y899D and F901D NRP1 tail peptides and binding was measured by ITC. Top panel shows the raw data and bottom panel shows the integrated and normalised data fitted with a 1:1 binding model. The kD values were measured over N = 3. **(L)** HEK293T cells were co-transfected to express a wild-type GFP-tagged NRP1 tail construct or mutants forms lacking the ESCPE-1 binding motif (ΔNYN) or the PDZ-binding motif (ΔSEA) and subjected to GFP-nanotrap. Endogenous ESCPE-1 subunits and GIPC1 were co-immunoprecipitated and band intensities were measured from N = 4 independent experiments. One-way ANOVA with Dunnett’s multiple comparisons tests. SNX6 ΔNYN p = 0.0019, SNX6 ΔSEA p = 0.9951, GIPC1 ΔNYN p = 0.6377, GIPC1 ΔSEA p = 0.0010. **(M)** Immunofluorescence staining and confocal microscopy of HeLa NRP1KO cells transfected with GFP-Nrp1 wt and GFP-Nrp1 ΔNYN. Non-permeabilised cells were labelled with anti-GFP, and signal intensity was quantified using Volocity software. Two-way ANOVA and Šídák’s multiple comparisons test. Total GFP: p = 0.9550; Surface aGFP: p = 0.7244. Scale bar = 50 µm. The bars, error bars and circles represent the mean, SEM and individual data points, respectively. ∗p< 0.05, ∗∗p< 0.01, ∗∗∗p< 0.001, ∗∗∗∗p< 0.0001.

**Extended Data Fig. 5.**
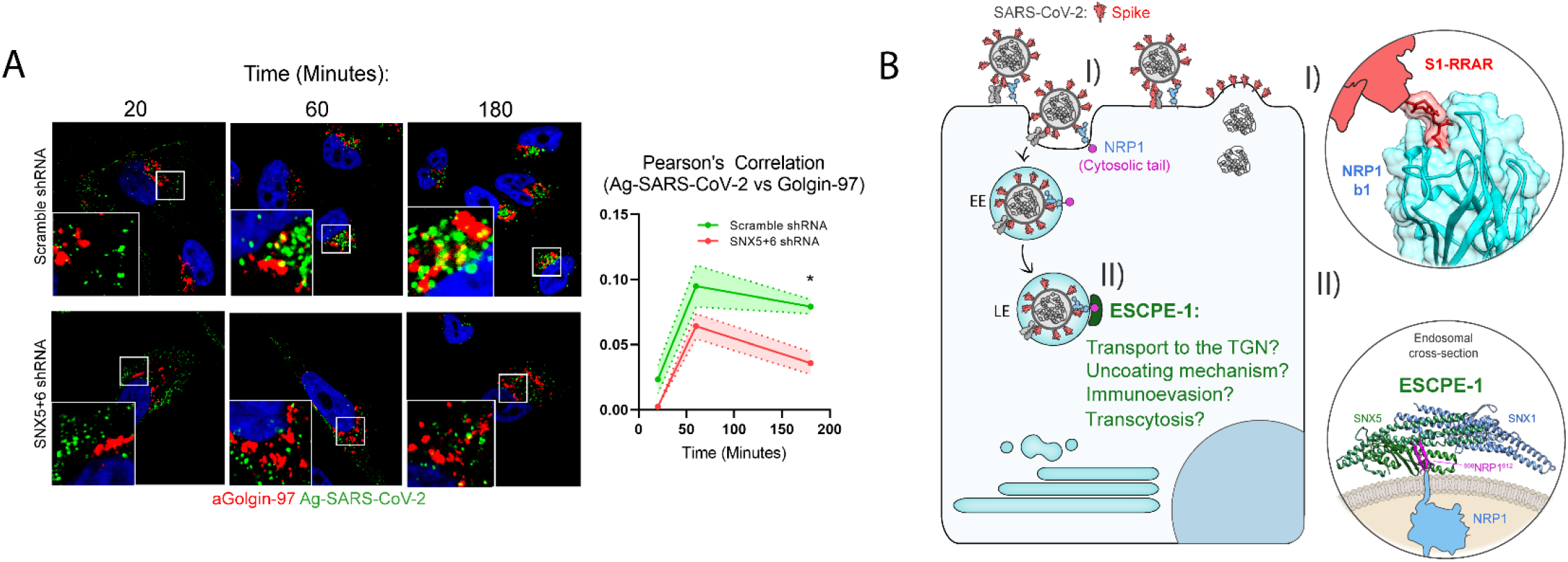
Model for ESCPE-1-Dependent Trafficking of Internalised NRP1-Bound SARS-CoV-2. **(A)** Ag-SARS-CoV-2 uptake assay in Scramble shRNA PPC-1 cells or SNX5+6 shRNA PPC-1 cells. Pearson’s correlation between Ag-SARS-CoV-2 and anti-Golgin-97 is measured over time using Image J software (N = 3 experiments). Two-way ANOVA with Šídák’s multiple comparisons test, p = 0.4040 (20 minutes), p = 0.1316 (60 minutes), p = 0.0259 (180 minutes). Scale = 20 µm**. (B)** Model for ESCPE-1-dependent trafficking of Internalised SARS-CoV-2. (I) SARS-CoV-2 engages NRP1 at the cell surface via direct binding between the CendR motif of S1 and the extracellular NRP1 b1 domain. (II) Once internalised into the endosomal network, the cytosolic tail of NRP1 is directly recognised by ESCPE-1, facilitating ESCPE-1 coat oligomerisation, membrane deformation and coupling to the cytoskeleton, along with potential consequences for SARS-CoV-2 biology.

**Supplementary Video 1**

HeLa cells were transfected with GFP-Nrp1. 24 hours after transfection, cells were live imaged using a confocal laser-scanning microscope at 37°C and incidences of GFP-Nrp1 (greyscale) localisation on tubular structures were observed. Representative frames are displayed in Extended Data Figure 3D.

**Supplementary Video 2**

HeLa cells were co-transfected with GFP-Nrp1 and mCherry-SNX1. 24 hours after transfection, cells were live imaged using a confocal laser-scanning microscope at 37°C and incidences of GFP-Nrp1 (green) and mCherry-SNX1 (red) colocalisation on tubular structures were observed. Representative frames are displayed in Figure 3D.

**Supplementary Video 3**

HeLa cells were co-transfected with GFP-Nrp1 and mCherry-SNX1. 24 hours after transfection, cells were live imaged using a confocal laser-scanning microscope at 37°C and incidences of GFP-Nrp1 (green) and mCherry-SNX1 (red) colocalisation on tubular structures were observed. Representative frames are displayed in Extended Data Figure 3E.

**Supplementary Table 1.** Complete list of proteins identified by SILAC-based proteomics following proximity labelling by HRP-TGN46. Untransfected HeLa cells were labelled in light (R0K0) SILAC media and treated with BP + H_2_O_2_, HRP-TGN46-expressing HeLa cells were labelled in medium (R6K4) SILAC media and treated with BP + H_2_O_2_, and HRP-TGN46-expressing HeLa cells were treated with BP in the absence of H_2_O_2_. N = 5 independent replicates. 10 proteins that were significantly enriched in the heavy (HRP-TGN46 expressing cells incubated with BP in the absence of H_2_O_2_) relative to untransfected HeLa cells, some of which were endogenous biotin-binding proteins, are highlighted in red and were removed from subsequent analyses.

**Supplementary Table 2.** Filtered list of proteins reproducibly enriched by HRP-TGN46 biotinylation. Proteins that were statistically significantly enriched in HRP-TGN46 expressing cells (medium condition) relative to untransfected HeLa cells (light condition) (p < 0.05, Log_2_ fold change >2), or present in ≥ 4 medium samples but < 3 light samples are presented.

**Supplementary Table 3.** Gene ontology analysis of enriched cellular components in the HRP-TGN46-labelled proteome. The list of proteins labelled by HRP-TGN46 **(Supplementary Table 2)** was analysed with PANTHER gene ontology software. The top 50 significantly enriched/depleted cellular component categories are displayed, ranked by p-value. + = significant overrepresentation, - = significant underrepresentation of categories.

**Supplementary Table 4.** Gene ontology analysis of enriched biological processes in the HRP-TGN46-labelled proteome. The list of proteins labelled by HRP-TGN46 **(Supplementary Table 2)** was analysed with PANTHER gene ontology software. The top 50 significantly enriched/depleted biological process categories are displayed, ranked by p-value. + = significant overrepresentation, - = significant underrepresentation of categories.

**Supplementary Table 5.** Enrichment of mannose-6-phosphate (M6P)-tagged proteins in the HRP-TGN46-labelled proteome. M6P-tagged proteins identified in ^17^ that were significantly enriched compared to untransfected HeLa cells (light condition) are displayed.

**Supplementary Table 6.** Complete list of proteins identified by SILAC-based proteomics following proximity labelling by HRP-TGN46 in Scr and SNX5+SNX6 siRNA-treated cells. Scramble siRNA-treated cells were labelled in light (R0K0) SILAC media, and SNX5+6 siRNA-treated cells were labelled in medium (R6K4) SILAC media, both conditions were biotinylated with BP and H_2_O_2_ and subjected to streptavidin affinity isolation. N = 4 independent replicates.

**Supplementary Table 7.** Filtered list of identified proteins from siRNA suppression screen. The list of proteins identified by SILAC-based proteomics in **Supplementary Table 6** was filtered to only include proteins present in the previously established HRP-TGN46-labelled proteome in **Supplementary Table 2**.

**Supplementary Table 8.** List of enriched and depleted proteins in the SNX5+6 siRNA HRP-TGN46-Labelled proteome. Proteins with a fold change of Log_2_ ± 0.26 and a p-value of < 0.1 are displayed. Proteins that were depleted from the SNX5+6 siRNA proteome relative to the Scr siRNA proteome are classified by the presence of a transmembrane domain, and putative cytosolic ΩxΦxΩ SNX5/6 interacting motifs. Proteins that were either identified as ESCPE-1 interactors^19^ or depleted from a GFP-SNX5(F136D) surface proteome^20^ are also indicated.

**Supplementary Table 9.** Gene ontology analysis of cellular component categories depleted in the SNX5+6 siRNA HRP-TGN46-labelled proteome. The list of proteins depleted in the SNX5+6 siRNA HRP-TGN46 proteome **(Supplementary Table 8)** were analysed with PANTHER gene ontology software. Significantly enriched/depleted biological process categories are displayed, ranked by p-value. + = significant overrepresentation, - = significant underrepresentation of categories.

**Supplementary Table 10.** Gene ontology analysis of biological process categories depleted in the SNX5+6 siRNA HRP-TGN46-labelled proteome. The list of proteins depleted in the SNX5+6 siRNA HRP-TGN46 proteome **(Supplementary Table 8)** were analysed with PANTHER gene ontology software. Significantly enriched/depleted biological process categories are displayed, ranked by p-value. + = significant overrepresentation, - = significant underrepresentation of categories.

**Supplementary Table 11.** Gene ontology analysis of molecular function categories depleted in the SNX5+6 siRNA HRP-TGN46-labelled proteome. The list of proteins depleted in the SNX5+6 siRNA HRP-TGN46 proteome **(Supplementary Table 8)** were analysed with PANTHER gene ontology software. Significantly enriched/depleted biological process categories are displayed, ranked by p-value. + = significant overrepresentation, - = significant underrepresentation of categories.

