## Supplementary material for "ESCPE-1 Mediates Retrograde Endosomal Sorting of the SARS-CoV-2 Host Factor Neuropilin-1": Molecular Modelling Methodology.pdf

### Supplementary Information

### Modelling the SNX5-NRP1 Complex

The aim was to generate any models of the NRP1 cytosolic tail sequence with a beta hairpin that could be mapped onto crystal structures of beta hairpins bound to SNX5 (pdb codes 6n5X, 6n5Y, 6n5Z).

Using the NRP1 cytosolic tail sequence:

*YCACWHNGMSERNLSALENYNFELVDGVKLKKDKLNTQSTYSEA.*

A sequence similarity search in the pdb[1] for regions of similarity, produced nothing. HHPRED[1] and Modeller[2] failed because there were no suitably similar homologous structures on which to build a model. The RAFT[3] software was used to generate folds ab initio, but this only produced all helix-coiled-helix structures. Itasser[4], which combines homology and ab initio methods, produced templates with some stretches of beta but none had a convincing hairpin.

Returning to the 6n5z structure in the pdb a Clustal Omega[5] alignment of the NRP1 residues region known to bind SNX-5 with those of the semaphorin-4C residues that bind to SNX-5 in the crystal structure 6n5z.pdb.

```
>NRP1_SNX5-binding-residues.
SALENYNFELVDGVKLKKDKLNTQ
>SEMA4C_ SNX5-binding-residues.
NWDPVGYYYSDGSLKIVP
CLUSTAL O(1.2.4) multiple sequence alignment
NRP1_bit. SALENYNFELVDGV-KLKKDKLNTQ 24
SEMA4C.   -NWDPVGYYYSDGSLKIVP       18
             +  +   **   +
```

Although not too promising, the hairpin DGs did align with the VDGV of NRP1 sequence which was predicted by iTasser to be part of a short stretch of beta secondary structure. The NRP1 corresponding residues were modelled along the SEMA-4C structure with the DG residues as the anchor point. The charged residues mapped well with the corresponding residues on SNX5.

The resulting NRP1 hairpin-SNX5 complex was energy minimised and subjected to molecular dynamics simulation using GROMACS[6] (2019.2) according to the previously published procedure[7]. 20 ns atomistic molecular dynamics allowed the backbone and side chain residues to relax. It was encouraging to see that the complex showed no signs of dissociating. To test this model further, residues upstream and downstream of this beta hairpin (according to the model generated with iTasser) were attached, energy-minimised and subjected to 20 ns molecular dynamics simulation as before. This extended hairpin peptide showed no signs of dissociating under these short simulation conditions. This conformation of the NRP1-SNX5 complex became one of the models proposed for experimental exploration.
